# Breast cancer cohort study identifies an effective, non-invasive, breath-based diagnostic linked to cellular environment-dependent, novel methylation metabolisms

**DOI:** 10.64898/2026.05.21.726956

**Authors:** Theo Issitt, Amy Harmens, Andrew S. Mason, James Turvill, Jenny Piper, Sean T. Sweeney, William J. Brackenbury, Kelly R Redeker

## Abstract

Volatile organic compounds (VOCs) demonstrate promise as non-invasive diagnostic tools. However, lack of mechanistically linked VOCs with biomarker discovery platforms limit delivery to the clinic. In this cohort-based study, we observed significant alterations of chloride-containing volatile fluxes, inclusive of methyl chloride (MeCl), in the breath of cancer patients that are consistent with our newly described metabolic model, derived through *in vitro* cellular assays and *in vivo* mice models. The diagnostic accuracy of this cohort study (60 patients) is equivalent to mammogram approaches. This newly identified and novel metabolism along with associated diagnostic biomarkers were initially identified through headspace studies of breast cancer cell lines which were deprived of serum, glucose or oxygen, similar to cellular conditions in tumors. In these cellular assays MeCl was consistently informative of cellular stress. Under resource limited conditions cellular production of MeCl was significantly reduced, and in several cases, cellular metabolism shifted to consumption. We present a new “push-pull” model in which cellular production of MeCl is linked to cellular methylation potential and methyl-transferase activity while consumption of MeCl is associated with methionine generation. Neither consumption nor production metabolisms have been described or quantified in humans or human tissues previously. The cellular headspace-derived model was tested using xenograft tumour bearing mice, which demonstrated reduced MeCl production, consistent with this model This work therefore presents a potentially powerful breath biomarker for cancer that translates from cellular and mice models through to human subjects.

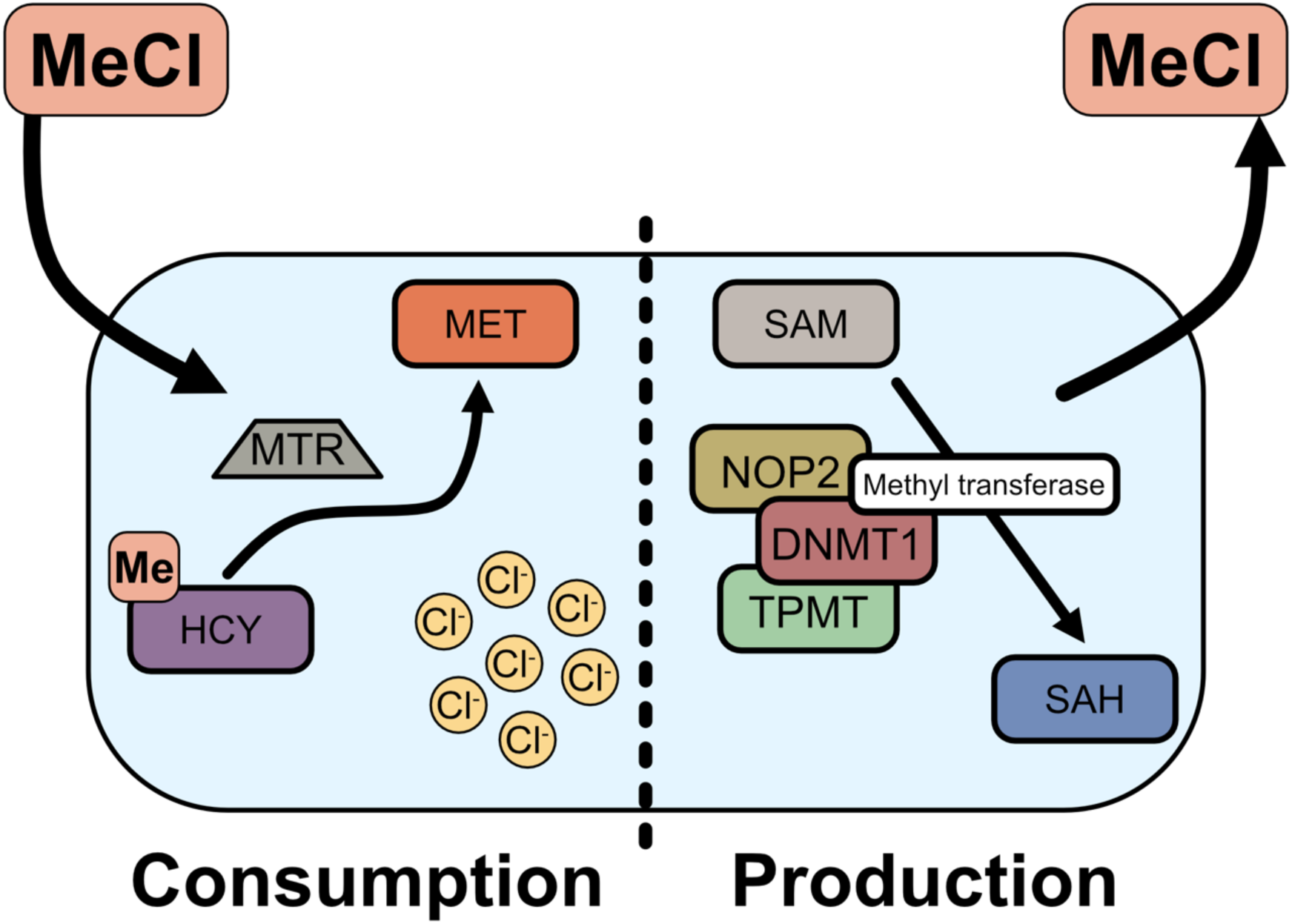

**Highlights:**

- (Poly)chloromethane compounds, including methyl chloride and chloroform, are indicative of cancer status in breath sampled from breast cancer clinic patients
- Glucose, serum and oxygen starvation induce significant changes in cellular metabolic fluxes
- Reduced methyl chloride production is indicative of cellular stress *in vitro*
- Methyl chloride production is linked to cellular methylation activity
- Methyl chloride consumption is linked to methionine synthesis, cellular enrichment in chloride concentration, and enhanced chloroform fluxes
- Methyl chloride fluxes in *in vitro* cell cultures under pathophysiologically relevant conditions translate to the breath of tumour bearing mice

## Introduction

Breath-based diagnostics have a number of benefits in provision of health services. Breath-based diagnostics are non-invasive, easily transported, and have the potential to aid in the delivery of personalized medicine. Consequently, many breath-based studies have been conducted and, while a few have been able to effectively diagnose disease in humans, our limited understanding of the biological and mechanistic processes driving these changes continues to reduce diagnostic efficacy [1]. Gaseous breath-based diagnostics rely on volatile metabolomics, which quantifies volatile organic compounds (VOCs), produced or consumed by biological processes in order to describe metabolic changes in organisms or systems. VOCs are small, carbon containing compounds which are gaseous at room temperature, often with a distinct odour, such ethanol or acetone.

Breath-based approaches have recently recognized that cellular environment plays a significant role in cellular metabolism. There have been a range of VOC studies using cellular models such as lung [2, 3], breast [4, 5] and liver [6] cancer, stem cells [7] and umbilical vein endothelial cells [8]. However, most studies to date have not effectively linked alterations in compounds associated with disease, such as cancer, to mechanistic processes driven by cell type. Our previous research has shown that variations in cell type (cancerous vs non cancerous) can be ascertained through their volatile profiles [5] and that cellular response to stress, from doxorubicin [5] or low oxygen [9], produced distinct and significant changes in cellular VOC profiles. Translation of VOC signals into more complex systems (i.e. the human body) introduces confounding and conflicting signals [1]. While cell type comparisons can be powerful diagnostic tools, they may be lacking in modelling the environmental pathological conditions *in vivo*. Throughout our previous work, translational characteristics in the VOC profile were more tightly determined by models in which cells experienced relevant resource conditions, *in situ* [5, 10]. From this we hypothesise that pathophysiologically-relevant systems are critical for effective VOC biomarker discovery.

The poor connection between VOCs and disease status in the published literature may be due to the influence of local environment on VOC metabolism in cellular models, since response to drugs [5], oxidative stress [11] and hypoxia [9, 12] have all shown that stress-dependent VOC metabolisms are linked to disease pathology. Host microbiome and infectious agents have also been shown to impact VOC metabolisms [13, 14], presumably through modification of resource and environment.

In this study we set out to link, or translate, the VOC profile of the aggressive, triple negative MDA-MB-231 breast cancer cell line to VOC profiles of mice bearing MDA-MB-231 xenograft tumours [15] and thence to human breast cancer patients. Initial quantification of VOC fluxes for MDA-MB-231 breast cancer cells and non-transformed MCF10a breast-derived cells under serum, glucose and hypoxic starvation (to reflect substrate starvation experienced by growing tumour cells; [15]) identified that reduced production (or even uptake) of methyl chloride was a consistently descriptive biomarker for cellular stress and, potentially, cancer [5, 9]. Tumor-bearing mice also demonstrate reduced production of methyl chloride, consistent with the physiologically-relevant cellular models [2].

MeCl is the most abundant atmospheric halocarbon compound (∼500pptv, [16]) and is commonly exhaled in human breath at substantially elevated concentrations relative to ambient (>1000 pptv) [17, 18]. In plants and fungi, MeCl is well known to be produced from s-adenosyl methionine (SAM) dependent methylation activity [19, 20] whereas in methylotrophic bacteria ambient atmospheric concentrations of MeCl are consumed [21]. We hypothesised that observed MeCl production is linked to SAM-dependent methylation activity. Methylation metabolisms were therefore investigated to explain observed variations in volatiles detected in *in vitro* cellular and *in vivo* mice models. We then tested whether cellular and mice-translatable outcomes were consistent in human patients through a clinical trial involving 60 patients that were being screened for breast cancer [18].

## Methods

### Orthotopic xenograft breast tumour model

Animal experiments have been reported in-line with the ARRIVE guidelines [22].Our operational tumour model has previously been described [23]. *Rag2^−/−^ Il2rg^−/−^* mice (bred in-house) were housed within temperature-controlled rooms, in individually ventilated cages with enrichment (3-5 mice per cage) and access to water and food ad libitum. At 6 weeks of age, female mice were anaesthetised (2% isoflurane in oxygen (2 L/min)) and 5 × 10^5^ MDA-MB-231 cells (suspended in Matrigel, 50% v/v in saline, 50 µL total volume) were injected into the left fourth inguinal mammary fat pad. Animal weight, condition and tumour growth (calliper measurement) were monitored daily. Mice were euthanized at 5 weeks after cell implantation or before tumours reached 15 mm diameter and tumours isolated. Mammary fat pad, without tumour, was also collected from the same animals (contralateral side).

We have previously published our method for sampling mouse headspace [5]. Nine-week-old female *Rag2^−/−^ Il2rg^−/−^* mice were selected for sampling. Experimental replicates were 2 mice from a cage across 3 separate litters/cages: 6 mice in total for each experimental group (control or with tumour as littermates). Mice were chosen randomly for experimental groups in each litter, blinding was not possible with animal care, sampling and analysis performed by TI.

Using tube handling methods, mice were gently placed, along with a cardboard tube and blue paper, into the custom-built chambers. Mice were allowed to acclimatise in undisturbed conditions during the initial 10 min chamber air equilibration period, accomplished by using a Yamitsu air pump with a flow rate of 750 mL per min. T0 samples were then taken and chambers were sealed for 20 min after which time T1 samples were taken.

At 4 weeks post implantation, volatile headspace samples were taken with the custom-built chambers, as described above, first with mice and then afterwards without the mice present but with the faecal material deposited during the initial sampling period. The number of faecal pellets were recorded once mice were removed from the chamber and was used to normalise VOCs against faecal VOCs (for comparison, and as shown in Figure S1).

### Human Patient breath sampling and flux analysis

Details of breath VOC pilot study design, patient selection criteria and information, breath collection and analysis have previously been reported [18].

Breath samples were collected at York and Scarborough NHS Trust (York Hospital) in a pilot study of 60 women referred through the “two-week wait” pathway for suspected breast cancer. Breath was collected using a novel dual-sampling platform that enabled concurrent collection of both inhaled and exhaled breath, which allowed volatile fluxes for each compound of interest to be quantified.

Patients provided 5 min duration breath samples while at rest through a custom sampling circuit with separate 5L stainless steel reservoirs for inhaled and exhaled breath. After the patient completed the 5 minute breathing period the 5L reservoirs were sealed and breath samples were transferred, via pressure differential, into 0.5L evacuated electropolished stainless-steel canisters (LabCommerce, San Jose, USA) for later analysis (maximum time between sampling and analysis < 1 week) by gas chromatography-mass spectrometry [24].

Systemic metabolic flux within the human body can be translated from these breath samples. Subtracting inhaled air content from exhalate air and multiplying by the average human breathing volume of ∼11,000L day^−1^ allows us to calculate metabolic flux within the individual in terms of µgrams compound individual^−1^ min^−1^.

This work presents data from patients ≥50 years old with grade 2/3 breast cancer compared to age-matched controls without cancer, focusing on (poly)chlorinated volatile organic compounds.

### Cell Culture and treatments

MDA-MB-231 and MCF7 breast cancer cells (a gift from Professor Mustafa Djamgoz, Imperial College London) were grown in Dulbecco’s Modified Eagle Medium (DMEM, Thermo Scientific, Waltham, MA, USA), 25 mM glucose, supplemented with L-glutamine (4 mM) and 5 % foetal bovine serum (Thermo Scientific, Waltham, MA, USA). The nontransformed human epithelial mammary cell line MCF10A (a gift from Prof. Norman Maitland) was grown in DMEM/F12 (Thermo Scientific, Waltham, MA, USA) supplemented with 5% FBS, 4 mM L-glutamine (Thermo Scientific, Waltham, MA, USA), 20 ng/mL EGF (Sigma-Aldrich, Roche; Mannheim, Germany), 0.5 mg/mL hydrocortisone (Sigma-Aldrich, Burlington, MA, USA), 100 ng/mL cholera toxin (Sigma-Aldrich, Burlington, MA, USA) and 10 µg/mL insulin (Sigma-Aldrich, Burlington, MA, USA). Cell culture media were supplemented with 0.1 mM NaI and 1 mM NaBr (to model physiological availability of iodine and bromide). Both formulations were considered control conditions. Serum free media and glucose free media were prepared in the same way but without those individual components. All cells were grown at 37 °C with 5 % CO_2_.

Prior to volatile collection, cells were trypsinised, and 500,000 cells were seeded into 8 mL complete media in 10 cm polystyrene cell culture dishes. Cells were then allowed to attach for 3-4 h, washed with warm PBS and 6 mL treatment media was applied. Volatile headspace sampling was performed 24 h later.

Cells were treated with 10 µM [25] 5-azacytidine (5-AZA, prepared in sterile water, Sigma-Aldrich, Germany) to block methylation events; the MAT2A inhibitor to block production of MAT (the enzyme which primarily catalyses the synthesis of S-adenosylmethionine), FIDAS-5 (10 µM, prepared in DMSO, Sigma-Aldrich, Germany) or sodium nitroprusside to block methionine synthase activity (SNP, 400 µM prepared in water, Sigma-Aldrich, Germany) where stated. S-adenosylmethionine (SAM) amendments were performed at a concentration of 50 µM, which has been determined to be sufficient to produce significant effects upon MDA-MB-231 without considerable cell death or cell growth [26, 27]. Drug concentrations were determined as producing a significant effect without excessive cell death over 24 hours, from the literature and with alamar blue assay; 5-AZA treatment was performed at 10 µM [25], FIDAS-5 at 10 µM [28], SNP at 400 µM [29].

### Cell culture VOC headspace collection and post-VOC cellular sampling and analysis

The method of headspace sampling has been described in detail [5, 9]. Briefly, cells placed in static polycarbonate chambers were flushed with lab air (4 L/min for 10 min) and time zero samples taken into pre-evacuated, electropolished stainless steel canisters (LabCommerce, San Jose, USA). Isolated headspace chamber samples were then left on a rocker for 120 min at the slowest setting, after which another canister sample was collected from the entrapped headspace. Cells were removed from the chamber, washed with PBS twice and lysed in 500 µL RIPA buffer (NaCl (5 M), 5 mL Tris-HCl (1 M, pH 8.0), 1 mL Nonidet P-40, 5 mL sodium deoxycholate (10 %), 1 mL SDS (10%)) with protease inhibitor (Sigma-Aldrich, Roche; Mannheim, Germany). Protein concentration of lysates was determined using BCA Assay (Thermofisher, USA). Background (medium only) readings were taken for all medium types following 24 h incubation at 37 °C and 5% CO_2_ (Figure S2A).

### GC/MS analysis of VOCs

All stainless steel VOC canister) samples (inclusive of cellular and mouse headspace samples as well as patient collected breath samples) were condensed onto a liquid nitrogen trap before being transferred, via heated helium flow, to an Agilent/HP 5972 MSD system (Santa Clara, CA, United States) equipped with a PoraBond Q column (25m x 0.32mm x 0.5 µm film thickness) (Restek©, Bellefonte, PN, United States), as previously detailed [5, 9]. Samples were analysed in selected ion monitoring (SIM) mode, (mice and human breath samples were also run separately in SCAN mode), specific details of which are available in Issitt et al. 2022 [5, 18].

Peak area/moles injected were calibrated from known standard injections. Moles injected for each compound of interest were quantified using these area-based calibration curves, and parts-per-trillion-by-volume (ppt) concentrations calculated by dividing moles of compound by moles of air injected (and multiplying by 1 trillion). Sample VOC concentrations were then normalised to CFC-11 concentrations (240 parts-per-trillion-by-volume (ppt)) [5, 10, 18, 30]. CFC was used as an internal standard as atmospheric concentrations of CFC-11 are globally consistent and stable and there are no known biological metabolisms in humans or mice that rapidly influence its ambient, atmospheric concentration [24]. Detailed equations are available [5, 9].

Where appropriate, media-only volatile fluxes (“backgrounds”) were subtracted from cellular plus media volatile fluxes and were then normalised to protein concentration. No variation between media types was observed (Figure S2) and so these were pooled to create an average media flux blank value per volatile. We have previously demonstrated no variation in DMSO containing media volatile flux for the compounds discussed in this research [5].

### MTT assay

MDA-MB-231 and MCF10A cells were seeded onto 96-well plates at a density of 8000 cells per well. Medium was changed once the cells had attached to the plate (4 h). Cells were then placed in cell culture incubation conditions described above. A total of 24 h later, 20 µL of MTT solution was added to each well and incubated for a further 3 h. Medium was removed, and precipitates solubilised in 100 µL DMSO. Absorbance was then measured at 570 nm using a Clariostar Plus microplate reader (BMG Labtech, Offenburg, Germany) across duplicate technical with triplicate experimental replicates.

### Sulphorhodamine B assay

To determine cell growth, sulphorhodamine B (SRB) assays were performed. The SRB assay measures cell density based on protein content [31]. Following incubation, cell monolayers were fixed with 10% (wt/vol) trichloroacetic acid (TCA) and stained for 30 min, after which the excess dye was removed by washing repeatedly with 1% (vol/vol) acetic acid. The protein-bound dye was dissolved in 10 mM Tris base solution for OD determination at 510 nm using a Clariostar (BMG) microplate reader [31] using duplicate technical with triplicate experimental replicates.

### Trypan blue exclusion assay

Trypan blue exclusion assay was performed on MDA-MB-231 and MCF10A cells following VOC sampling. Following a published protocol [32], trypsinised cells were mixed with 0.4% Trypan blue solution and counted to determine the number of unstained (viable) and stained (nonviable) cells. Cells were counted at 20x magnification across 10 fields of view using duplicate technical and triplicate experimental repeats.

### Ion Chromatography

A previously published protocol was adapted to quantify chloride contents of cells and tissues [33]. Cells were grown to 80% confluency in a 10 cm dish and treated for 24 h, as appropriate. Cells were washed in 10 mL pH 7 phosphate buffer twice and lifted using a cell scraper in 5 mL phosphate buffer. Cells were then pelleted by centrifugation (200 g for 5 min), supernatant aspirated and resuspended in 1 mL H_2_O. The pellet solution was then snap frozen in liquid nitrogen, thawed and pulse sonicated (5 s pulse for 20 s). A sample of 50 µL was removed at this point for protein quantification using BCA assay (Pierce, Thermo Scientific). The pulse sonicated mixture was then added to 5 ml H_2_O and syringe filtered through a 0.2 µm prerinsed (with ddH_2_O) PTFE filter. Tumour and mammary fat pad samples were weighed, washed in phosphate buffer, snap frozen, thawed and sonicated and treated as above.

Ion chromatography was performed using a Dionex ICS-2000 ion chromatograph fitted with an EGC III KOH Eluent generator cartridge, ADRS 600 2 mm suppressor, DS6 heated conductivity cell and AS40 autosampler. Dionex IonPac AS18 (2 mm i.d. x 250 mm length) analytical column fitted with an IonPac AG18 (2 mm i.d x 50 mm length) guard. Mobile phase gradient: 2 mM potassium hydroxide (hold 1 min) to 41 mM over 35 min (hold 4 min) at a flow rate of 0.25 ml/min column. A suppressor current of 26 mA, column temperature of 30 °C, detector temperature of 35 °C and a sample injection volume of 15 µL were used. Instrument control and data processing were performed using Chromeleon software. Samples were quantified against calibration curves derived from known standard injections.

### Methylation ELISA

All samples were tested for global DNA methylation using the MethylFlash™ Methylated DNA Quantification Kit (Colorimetric) (Epigentek, Farmingdale, NY, USA). MethylFlash uses an enzyme-linked immunosorbent assay (ELISA) based method to quantify global DNA methylation. For each sample, 200 ng of DNA was used, in duplicate, as recommended by Epigentek. DNA was isolated from cell pellets or tissue using a tissue and blood specific DNA extraction kit (Qiagen). The ELISA was performed following manufacturer’s instructions. Absorbance readings from each plate were measured at 450 nm using a Clariostar (BMG) plate reader and Mars software (BMG) across duplicate technical with triplicate experimental replicates.

### HPLC for methionine, homocysteine, vitamin B12, SAH, and SAM

Cells were grown to 80% confluence in 6 well plates and were then washed with PBS (5 mL x 2) and stored, sealed at −80 °C. Samples were prepared in ice cold 90% methanol with 0.1% trifluoroacetic acid (TFA) [34]. The following authentic standards were used: L-methionine (Met), L-homocysteine (Hom), cyanocob(III)alamin (B12), S-(5′-adenosyl)-L-homocysteine (SAH) and S-(5′-Adenosyl)-L-methionine chloride dihydrochloride (SAM), all supplied by Merck.

Standard stock solutions were prepared in water with 0.1% TFA at the following concentrations: Met 4 mg/mL, HCY 5 mg/mL, B12 6 mg/mL, SAH 4 mg/mL, SAM 2 mg/mL, and stored at −20°C for up to two weeks. Eight-level standard curves were constructed in 90% methanol at the following concentration ranges: Met 2.1 ng/mL-40 µg/mL, HCY 24 ng/mL-100 µg/mL, B12 38 ng/mL-2.4 µg/mL, SAH 9.8 ng/mL-40 µg/mL, SAM 0.31-20 µg/mL (the lower-end concentration for each compound was that at which a peak was clearly visible, and serves as an approximation of LOD).

Samples were analyzed with a Waters Acquity IClass UPLC, interfaced to a Waters Synapt G2-Si, operated in positive ESI sensibility mode (scan rate 0.6 s, mass range 50-1600 *m/z*). Mobile phase A) was 10 mM ammonium formate, pH 3.0 (adjusted with ammonium hydroxide) and 1% acetonitrile, mobile phase B) was acetonitrile. The gradient started at 90% B and decreased to 35% B over 19 min, where it remained until 24 min, to return to 90% B at 25 min and to re-equilibrate until 35 min. The flow rate was 0.5 mL/min. A MilliporeSigma SeQuant® ZIC-HILIC column (100 x 4.6 mm, 200 Å, 5 µm) was used at 40°C. The autosampler was kept at 7°C, injection volume was 10 µL. Analytes were identified according to accurate mass (all [M+H]+ adducts) and retention time, compared to authentic standards using Skyline software v22.2.0.351. Samples were stored at −80°C and loaded into the autosampler 20 min before each run as SAM degrades rapidly at room temperature [35].

### RNA sequencing analysis

Public RNA sequencing data was available for MCF10A and MDA-MB-231 cell lines (SRA:PRJNA302668; [36]), as well as MDA-MB-231 cells grown in normoxic and hypoxic conditions (1% O_2_) from either total RNA sequencing after ribodepletion (SRA: PRJNA604033; [37]) or mRNA sequencing after polyA enrichment (SRA: PRJNA530760; [38]). Following standard quality control, including adapter removal and removal of low-quality reads using Trimmomatic v0.36 [39], gene-level expression values in transcripts per million (TPM) were derived against the Gencode v41 human transcriptome using kallisto v0.46.1 [40]. TPMs were recalculated after exclusion of non-coding and mitochondrial genes. Differentially expressed genes in each experimental condition were identified using sleuth v0.30.0 [41] in R studio. MDA-MB-231 hypoxia datasets were also combined to provide a higher sample number, controlling for experimental design as a batch effect. Between experiment comparisons were drawn using π-values [42], a product of fold change and significance value. Methyl transferases were selected using identified methyl transferases [43] (Supplementary Table 1). π-values were used to direct a ‘prerank’ gene set enrichment analysis using a curated list of 198 methyltransferase genes (Supplementary Table 1) using GSEApy v1.0.5 [44].

### Transfection

Cells were transfected using the transfection reagent TransIT-siQuest (Mirusbio reagents) in OPTIMEM (Gibco). Scrambled (control) siRNA and siDNMT (SMARTpool) were purchased from ThermoFisher and used at a concentration of 25 nM with cells grown to 80% confluency, following the manufacturer’s instructions. Knockdown was confirmed by Western blot.

### Western blot

Cells were prepared for immunoblotting by lysing in RIPA buffer with phosphatase and protease inhibitor cocktails (Sigma-Aldrich, UK). 35 µg of protein lysate, determined by BCA assay (Pierce, Thermo Scientific), was resuspended in Laemmli buffer, denatured for 5 min at 95°C, separated by SDS PAGE and transferred to nitrocellulose membrane (Whatman, USA). Nitrocellulose membranes were immunoblotted with the following primary antibodies: rabbit polycolonal antihuman DNMT1 (Novus Biologicals, UK), mouse monoclonal antihuman MAT I/II (F-12, Santa Cruz Biotechnology, USA), mouse monoclonal antihuman PRMT1 (B2, Santa Cruz Biotechnology, USA), mouse monoclonal antihuman α-tublulin (Cell signalling, USA). HRP-conjugated secondary antibodies were used for chemiluminescence detection with Luminol (Santa Cruz Biotechnology, USA) and protein levels were quantified by densitometry with ImageJ (NIH, Bethesda US).

### Hydrogen peroxide (amplex red) assay

Experiments were performed in phenol red free DMEM. DMEM containing 50 μM Amplex Red reagent (Thermo Scientific, Waltham, MA, United States) and 0.1 U/mL horseradish peroxidase (HRP, Thermo Scientific, Waltham, MA, United States) was added to cells in 12 well plates (500 μL per well) for 15 min following 24 h in starvation, starvation plus SAM or control conditions. We have previously reported no change in ROS using this assay following 24 hours of hypoxia [9]. Fluorescence at 590 nm was measured with a plate reader (Clariostar, BMG, Ortenberg, Germany) and compared against a H_2_O_2_ standard curve for quantification.

### Database searching and alignment

The methyl chloride transferase (*Batis maritima,* UNIPROT ID: Q9ZSZ7) and cmuA methyltransferase (UNIPROT ID: F8J7J8 [45]) were aligned against the human proteome using Clustal (1.2.4) multiple sequence alignment by UNIPROT BLAST.

### Data analysis

Graphs and statistics were generated/performed in Graphpad Prism. Details of specific tests are provided in figure legends. RNA seq data were arranged in R studio following analysis (described above).

### Ethical approval

Approval for all animal procedures was granted by the University of York Animal Welfare and Ethical Review Body. All procedures were carried out under authority of a UK Home Office Project Licence and associated Personal Licences.

Breath sampling was performed at the Magnolia Centre at York hospital (York and Scarborough NHS Trust). The study received ethical approval from an NHS Research Ethics Committee (IRAS ID: 318636) and University of York Biology Ethics Committee (reference: KR202302). All participants provided written informed consent before any study procedures were undertaken. The study adhered to the ethical principles outlined in the Declaration of Helsinki and Good Clinical Practice guidelines. Approval covered patient recruitment, breath collection, data handling, and analysis as described in the protocol.

## Results

Our prior research has shown that production of methyl chloride is significantly reduced in the headspace of stressed (750 nM doxorubicin treatment) MDA-MB-231 cells relative to unstressed (no doxorubicin treatment) MDA-MB-231 cells[5]. Similarly, stress induced by hypoxia in MDA-MB-231 cells led to significant reduction in methyl chloride production [30]. Our meta-analysis of breath-based cancer diagnostics highlighted that a key limitation across studies was the lack of mechanistic understanding linking VOC biomarkers to underlying pathophysiology [46]. Furthermore, halogenated volatiles, including chlorinated and brominated compounds, have not been systematically investigated under pathophysiologically-relevant conditions that reflect the tumor microenvironment.

The tumor microenvironment can be considered a “stressful” environment, in that the local environment is more acidic, has greater interstitial pressure and is hypoxic. Tumor function under stress, combined with the well established enhanced permeability of tumor blood vessels suggests that the cellular impact of the stress induced by the cellular microenvironment induced by cancer should diffuse readily to the blood stream, and thereby into the lungs, where it can be detected in the breath of the patient [1].

### Volatile flux is altered in the breath of MDA-MB-231 tumour xenograft mice

The breath of mice with and without tumour xenografts was investigated to determine if VOC profiles of MDA-MB-231 cells would transfer to the breath of mice. Mice were age and sex matched, with consistent time point sampling at 4 weeks post tumour-induction protocol. Behaviour of two VOCs were significantly altered (Figure 1). MeCl flux was reduced (Figure 1C) and butanone was increased (Figure 5C) in tumour-bearing compared with control mice. No variations were seen in the VOCs from faecal material in mice with either treatment (Figure S1A).

**Figure 1.**
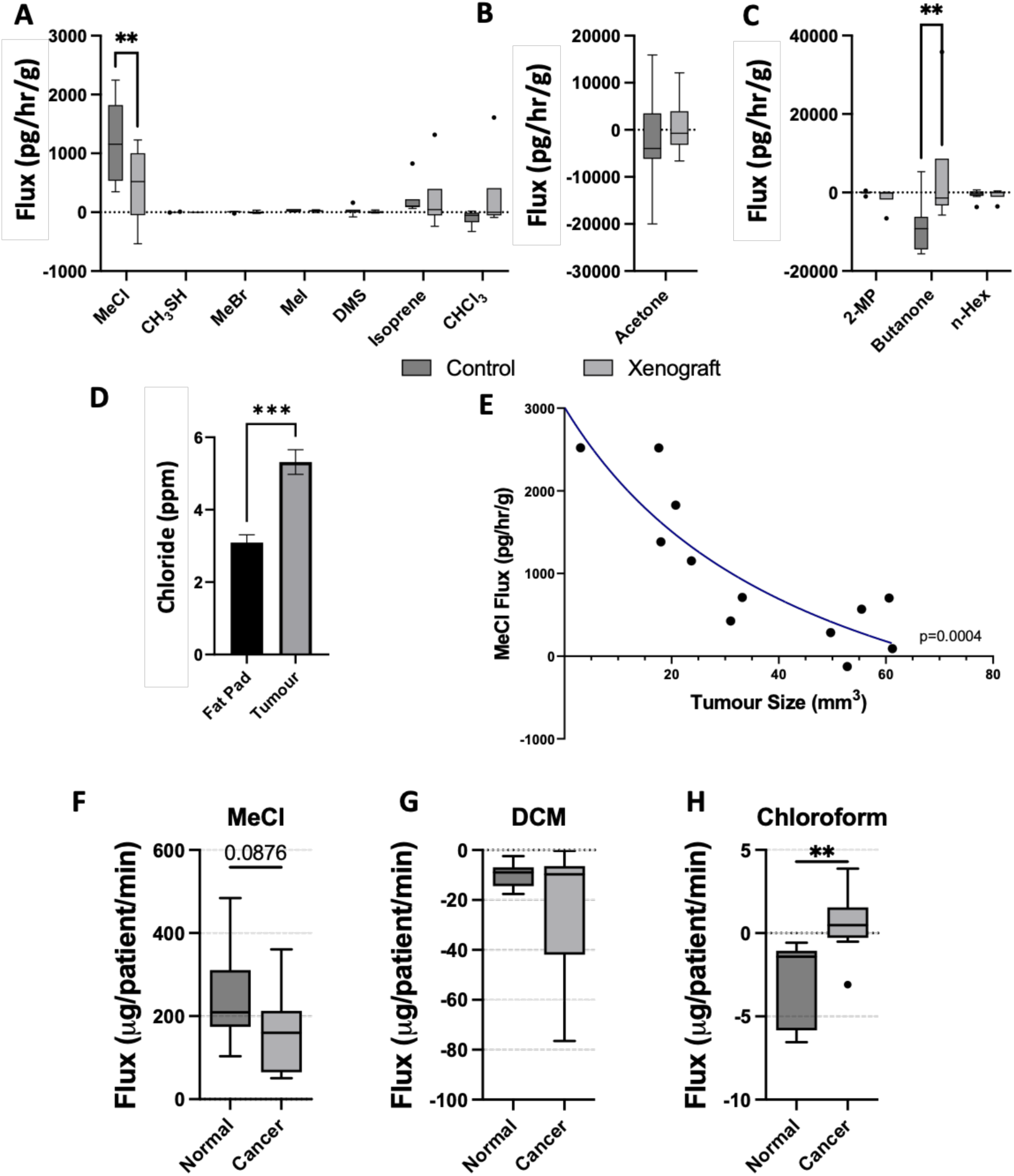
Volatile flux is altered in the breath of MDA-MB-231 tumour xenograft mice and breast cancer patients. **(A-C)** Volatile flux of mice breath in control and MDA-MB-231 xenograft tumour bearing mice. Faecal VOC fluxes were subtracted from mouse breath fluxes and residual fluxes normalised to mouse weight (pg/hr/g). **(D)** Intra-tumour and contralateral mammary fat chloride content in ppm (g Cl/g biomass). **(E)** Methyl chloride (MeCl) flux (pg/hr/g) of mouse breath (faecal MeCl flux subtracted and mouse weight normalised) for tumours of varying size (mm^3^). **(F-H)** (Poly)chloromethane compound volatile flux in the breath of over 50s women with grade 2/3 breast cancer versus normal patients (n=8 normal, 12 cancer). CHCI_3_ = Chloroform, DMS = Dimethyl sulphide, MeBr = Methyl bromide, MeI = Methyl iodide, CH_3_SH = Methanethiol. Boxplot whiskers show median ± Tukey distribution (n=6 for A-C, n= 20 for normal and 19 for cancer for F-H). Bar plot **(D)** shown with mean ± SEM. Two-way ANOVA with Bonferroni post hoc test was performed for **A** and **C**. Students T-test performed for **D, F-H**. Pade (1,1) approximant, least squares fit with non-linear fit test performed, shown by line of fit in **E**.

Chloride was investigated in tumours compared to the mammary fat pad without xenograft. Consumption, or reduced levels of MeCl production, significantly correlates with intracellular chloride accumulation in tumor biomass relative to contralateral mammary fat pad (from 3 ppm to 5 ppm, Figure 1D).

Tumour size was compared to MeCl flux, which revealed MeCl fluxes reducing as tumour size increased (Figure 1E). A non-linear regression with padé fit, showed this trend to be significant (p=0.0004). Chloride content of these tumours were investigated, however no significant trends were observed between tumor size and chloride content (Figure S1D).

### Volatile flux is altered in the breath of breast cancer patients

Breath volatile flux of breast cancer patients collected in a previous pilot study was investigated to test translation of outcomes to human subjects [18]. This work identified a significant variation in breath volatile flux linked to age and, once age groups were distinguished, was able to separate patients ≥ 50 years old with cancer or benign tumours from healthy volunteers. The data presented here is from those patients ≥ 50 with grade 2 or 3 breast cancer vs age-matched controls. Significant differences between tumour grades were observed [18]. Grade 1 tumours revealed significant differences in Chloroform and DMA flux. Furthermore,grade 2/3 tumours show increased levels of tissue abnormality, which is linked to increased levels of tumour stress [47]. Only grade 2/3 tumours are included here, in the context of this study investigating the role of pathophysiological stress of the TME.

MeCl in the breath of cancer patients was reduced compared with non-cancerous patients (Figure 1F, p = <0.1). There was no observable change in DCM (CH_2_Cl_2_; Figure 1G) however chloroform flux (CHCl_3_) was significantly increased in the breath of cancer patients compared with non cancerous patients (Figure 1H). Chlorine-containing biomarkers were compared to test translation of the VOC fluxes described here between pathophysiologically consistent cells, and mice and humans with tumours. Despite the disparity in cancer biomass to patient weight between humans and mice, we observe similar trends in MeCl and a substantive to significant increase in chloroform (Figures 1A and 1H).

### Cancer and non cancer breast cells produce different pathophysiological responses dependent upon starvation stress type

Hypoxia, pH and interstitial pressure are not the only stress inducing parameters within the tumour microenvironment. We further explored the relevance of pathophysiologically appropriate conditions on cellular function in cells by examining the implications of reduced energy substrate availability in cancer (MDA-MB-231) and non cancer (MCF10a) cells. Starving cells of serum and glucose models resource limitations as expected in tumour pathophysiology, since growing tumours experience poor availability of substrate delivery [48].

Overall, cellular response to glucose and serum starvation was similar between MCF10a and MDA-MB-231. In both cell lines MeCl and acetone fluxes became more consumptive while production of isoprene, butanone and n-hexane were enhanced under starvation-specific conditions. However, cancer and non cancer cells in starvation medium over 24 hours exhibited consistent and significant changes in flux for only 3 out of our 40 targeted compound treatments (i.e. – methyl chloride under serum and glucose treatments for MDA-MB-231 cells, would be equivalent to two compound treatments).

The conditions and compounds under which non cancer and.cancer cells behaved similarly included lack of serum treatments leading to significantly reduced MeCl production (often causing MeCl consumption) (Figure 2A, 2D), and lack of glucose treatments enhanced acetone consumption (Figure 2B, 2E) while production of butanone increased (Figure 2C, 2F). Broadly however, both breast cell types reduced MeCl production under starvation conditions, but non cancer cells showed a substantially reduced, often insignificant effect. Similarly, acetone consumption and butanone production increased for both cell types when under resource stress but these effects were inconsistent between starvation treatments.

**Figure 2.**
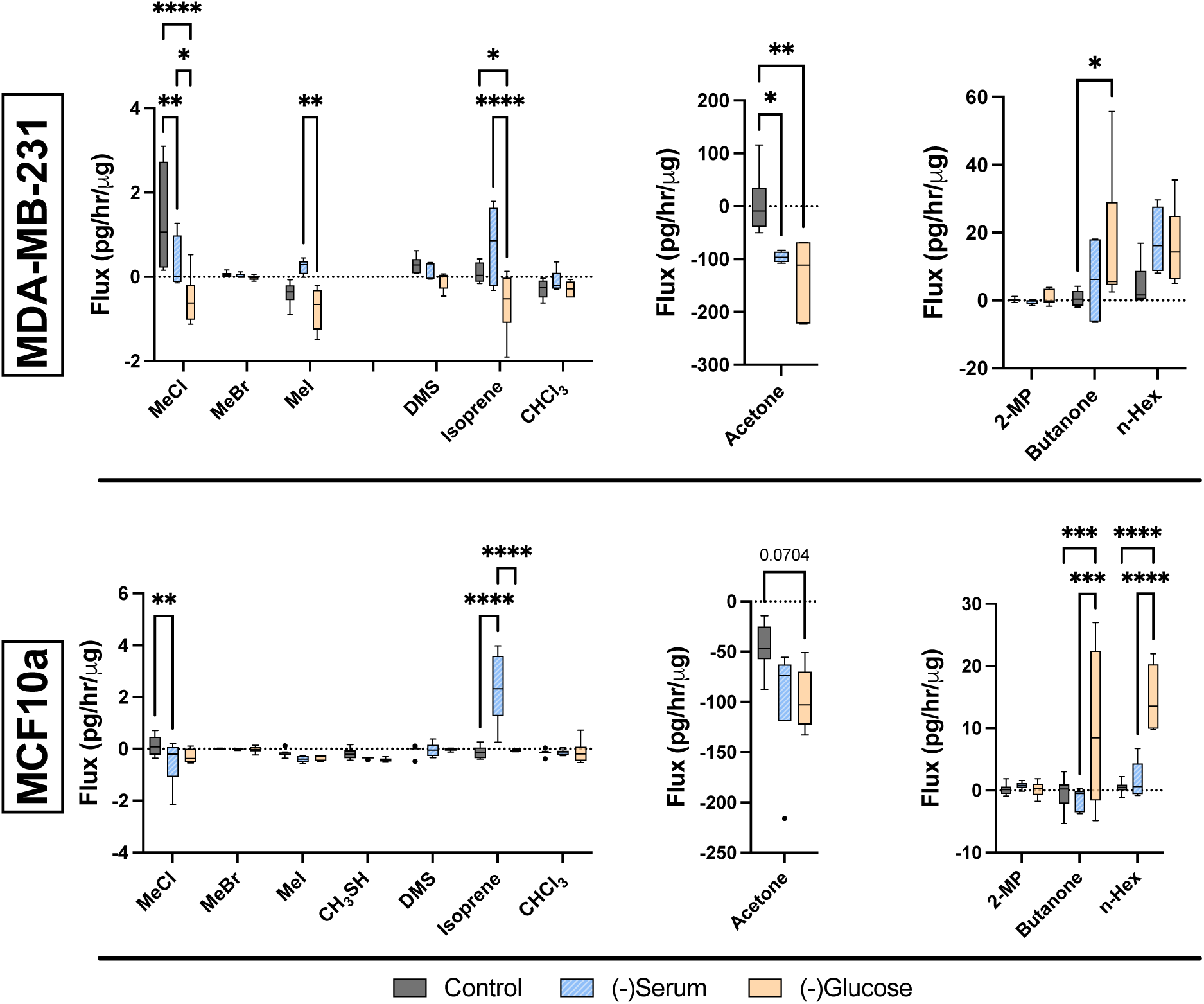
Starvation of breast derived cells produces detectable changes in select volatile organic compounds. Volatile flux (pg/hr/µg) for the cancerous MDA-MB-231 (**A-C**) and noncancerous MCF10a **(D-F).** Media subtracted and protein-normalised. (-)Serum = media without serum, (-)Glucose = media without glucose. MeCl = Methyl chloride, MeBr = Methyl bromide, MeI = Methyl iodide, CH_3_SH = Methanethiol, DMS = Dimethyl sulphide, CHCI_3_ = Chloroform, 2-MP = 2-methyl pentane, n-Hex = n-hexane. Boxplot whiskers show median ± Tukey distribution (n=6). Two-way ANOVA with Bonferroni post hoc test was performed for **A, C, D** and **F**. One-way ANOVA with Tukey post hoc test was performed for **B** and **E**; * *p* < 0.05; ** *p* < 0.01; *** *p* < 0.001; **** *p* < 0.0001.

Compounds which showed significant effects that were distinct to cancer cell type include: isoprene, for which glucose starvation caused significantly increased consumption in MDA-MB-231 cells whereas in non cancer cells serum starvation caused significantly enhanced production (Figure 2A, 2D). n-Hexane showed enhanced production under both starvation treatments and for both cell lines. However only MCF10a showed significant changes in measured fluxes (Figure 2C, 2F). Serum starvation produced an increase in MeI, and these fluxes were significantly different from glucose starvation (Figure 2A).

Glucose starvation of MDA-MB-231 cells produced a significant metabolic transformation, from production of MeCl to consumption (Figure 2A). Consumption of isoprene and acetone was also observed in glucose starved cells (Figures 2A and B), both of which deviated significantly from an effectively non-metabolised state (zero flux) when not starved.

Growth and metabolic activity of both cell lines under starvation was investigated to determine cellular responses to these stresses over time. At 24 hours, there was a considerable reduction in cells following serum starvation (Figure S2B and C). Glucose starvation produced no changes in growth for MDA-MB-231 cells and an increase in MCF10a over 24 hours, with relatively less growth after 48 hours compared to control (Figure S2B and C). Metabolic activity of cells was measured using MTT assay (Figure S2D and E). A reduction in metabolic activity was observed in all starvation conditions for both cell lines however the only significant difference was recorded for glucose starved MCF10a relative to control (Figure S2D). Starvation of MDA-MB-231 cells showed reduction in viability by MTT, without significance (Figure S2E). Cell death in these conditions was investigated using trypan blue assay. Both cell lines showed significant reduction in percentage of viable cells (between 50 and 70% cell death) in both serum and glucose free medium (Figure S2F and G).

### A note on the challenges inherent in complex cellular metabolisms

Fluxes of materials within cells can be modelled, or visualized, similarly to a box model as used in biogeochemistry[49]. In a cell-representing box model the change in the amount of any given substance within the cell over time can be represented through an equation similar to that below (using methyl chloride as an example):

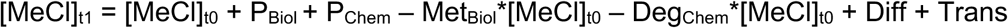

Where [MeCl] is the cellular concentration of methyl chloride, P_Biol_ and P_Chem_ are the production from time 0 to time 1 of methyl chloride by biological and chemical processes within the cell, Met_Biol_*[MeCl]_t0_ is the utilization of MeCl via enzymatic processes, Deg_Chem_*[MeCl]_t0_ represents the degradation of MeCl through chemical processes, and both which are generally dependent upon the concentration of MeCl in the cell, to a first order approximation. Diffusion of material in and out of the cell, as well as active transport of materials in an out of the cell are represented, per unit time step, as Diff and Trans.

This conceptualization of flux is challenging, since any change in flux, or change with time, may be due to multiple causes. Where we describe fluxes that may have decreased or increased under specific treatments, these observed changes can be explained in several ways. To simplify the equation above, we can assume that chemical production of methyl chloride in the body is much slower than any biological production, removing P_Chem_ from the equation. Likewise degradation of MeCl at human temperatures and conditions is likely to be much smaller than biological metabolism. If we also assume that diffusion and active transport are constant, then these terms can also be ignored in this context.

However, this still allows the potential that an increase in flux may be due to either i) an increase in biological production or a reduction in biological metabolism. Similarly, a reduction of flux may be due either to a decrease in production or an increase in utilization. A fully negative flux therefore does not require that the only active process is metabolic uptake, only that metabolic uptake dominates biological production.

### Methyl chloride flux positively correlates with chloride content and methylation rates in non stressed breast cells

MeCl production is significantly greater in unstressed MDA-MB-231 compared with MCF10a and MCF7 (Figure 3A and Issitt et al. 2022 [5]). Intracellular chloride was investigated using ion chromatography to determine whether reaction kinetics, driven by intracellular chloride concentrations, could partially explain MeCl flux changes. Intracellular chloride content showed a similar trend to MeCl flux within these cell lines; both intracellular chloride and MeCl fluxes were significantly greater in MCF7 and MDA-MB-231 cells when compared to MCF10a (Figure 3B).

**Figure 3.**
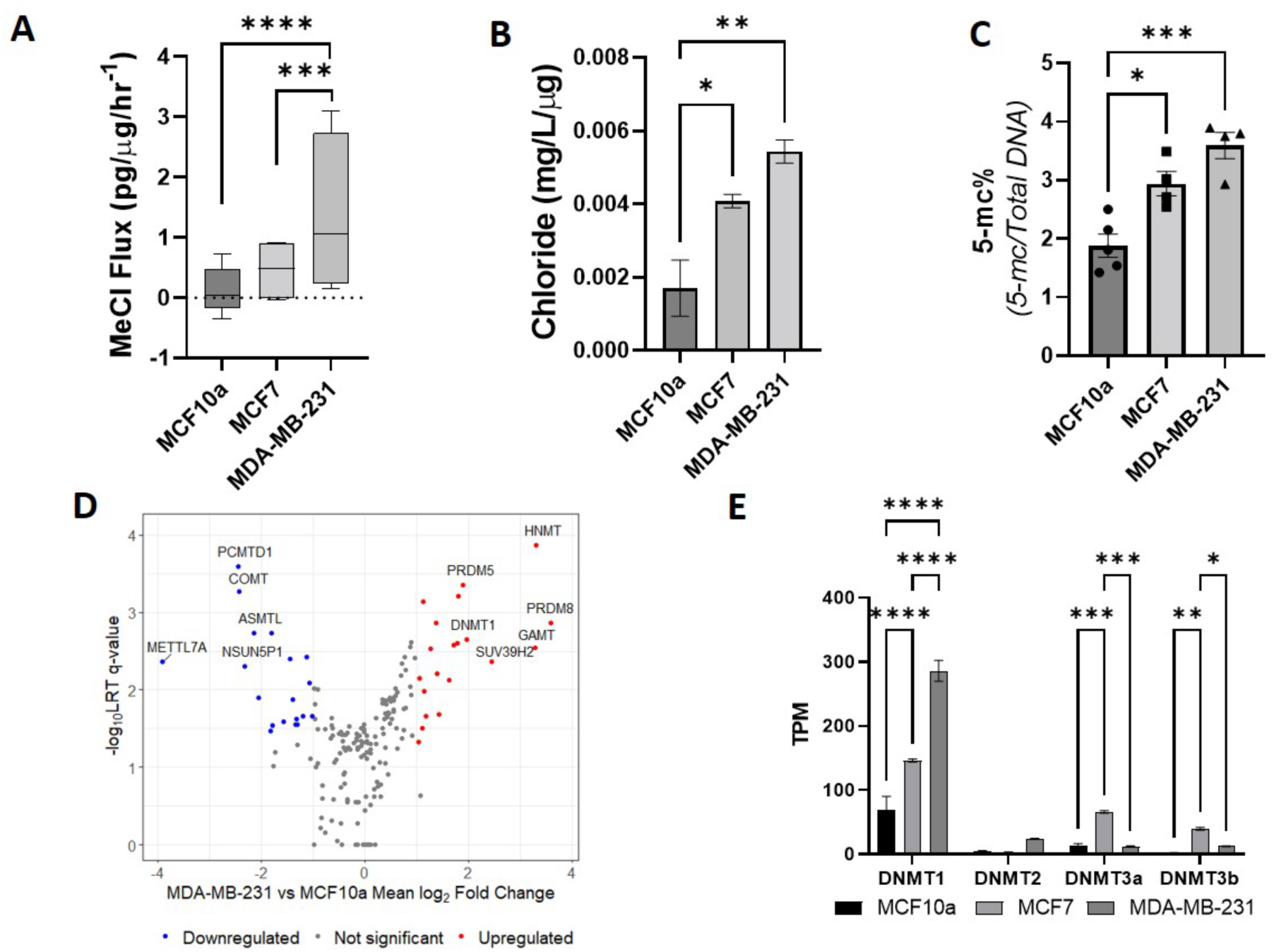
In non stressed cells methyl chloride flux corresponds with cellular chloride content and DNA methylation in breast derived cells. **(A)** Methyl chloride (MeCl) flux in pg MeCl/μg protein/hr, statistical analysis performed within expanded volatiles data set; two-way ANOVA with bonferonni post hoc test represented here independently (n=6). **(B)** Cellular chloride content in parts per million (ppm; mg/L/µg total protein) for different cell types. **(C)** Percentage of 5-methylcytosine (5-mC) of total DNA loaded for each cell type. **(D)** Volcano plot of RNA seq data for −log_10_ LRT (likelihood ratio test) q values (corrected p values using Benjamini-Hochberg) of methyl transferase genes shown vs mean log_2_ fold change of MDA-MB-231 vs MCF10a **(E)** Transcripts per million (TPM) of RNA seq data for DNA methyl transferases (DNMT) for different cell lines. Bar plots shown with mean ± SEM for **A-C** and mean ± SD for **E**. One-way ANOVA with Tukey post hoc test was performed for **B, C** and **E**; * *p* < 0.05; ** *p* < 0.01; *** *p* < 0.001; **** *p* < 0.0001.

Methyl chloride is known to be generated in other eukaryotic organisms through enzyme mediated transfer of methyl groups from s-adenosyl methionine (SAM) [19, 50, 51]. Enzyme-driven control of MeCl fluxes was therefore next considered. DNA methylation content of the three cell lines was investigated and compared with MeCl fluxes. 5-methyl cytosine residue percentage, determined by ELISA, of total DNA content within cells was seen to significantly increase from 1.8% for MCF10a, to 3% for MCF7 and 3.6% for MDA-MB-231 cells in control, unstressed conditions (Figure 3C).

There are no currently published or identified pathways for MeCl metabolism in humans, despite reports of its elevated concentration in human breath (>5x ambient) and interaction with the human gut microbiome [16]. MeCl production has been linked to s-adenosyl methionine dependent methylation in plants [19] and Maritime seawort [50, 51], therefore a protein:protein alignment was performed with BLAST from UNIPROT [52] for MeCl transferase (*Batis maritima,* UNIPROT ID: Q9ZSZ7) against the human database. The top result was thiopurine methyltransferase (TPMT, UNIPROT ID P51580), which was subsequently aligned, producing 24.8% similarity (Figure S3A). The alignment suggests similarity between proteins, with some conservation in the binding domain of TPMT (Figure S3A) providing a potential avenue for MeCl interaction.

Plant studies suggest multiple methyl transferases (MTs) may be involved in MeCl production [53]. RNA sequencing data was therefore investigated to identify variation in the TPMT gene and MTs generally. Methyl transferase RNA variation is shown as a volcano plot (Figure S3B). Differential expression analysis (Figure 3D) revealed 20 (red) and 19 (blue) MT genes (from a report list in Petrossian & Clarke, 2011 [43]) which were respectively up- and down-regulated in MDA-MB-231 cells. Of these, the six most significantly upregulated were *HNMT, PRDM8, SUV39H2, DNMT1, PRDM8*. The six most significantly upregulated MTs in MCF10a were *METTL7A, PCMTD1, COMT, NSUN5P1, ASMTL and COMTD1.* Whilst individual methyl-transferase gene expression differed between these divergent cell lines, gene seat enrichment analysis (GSEA) found no significant shift across the entire gene family (NES (normalised enrichment score): 0.971; FDR (false discovery rate): 0.503, Figure S4B).

To further describe observable increases in DNA methylation seen between cell types, DNA methylation specific genes were investigated as transcript per million (TPM) from the same public dataset. DNMT1 was increased in MCF7 and MDA-MB-231 compared to MCF10a, matching the changes observed in DNA methylation (Figure 3E) and consistent with changes in MeCl fluxes and intracellular chloride content. DNMT2 was slightly increased for MDA-MB-231 compared to other cells while MCF7 showed increases in DNMT3a/b (Figure 3E).

### In resource starved MDA-MB-231 cells methyl chloride flux positively correlates with methylation metabolism and inversely correlates with intracellular chloride content

We have demonstrated reduced MeCl production in response to serum starvation, and MeCl consumption during glucose (Figure 2A) and oxygen starvation [9] (consolidated in Figure 4A). Movement of chloride into and out of the cell also appears to be linked to cellular environment and stress.

**Figure 4.**
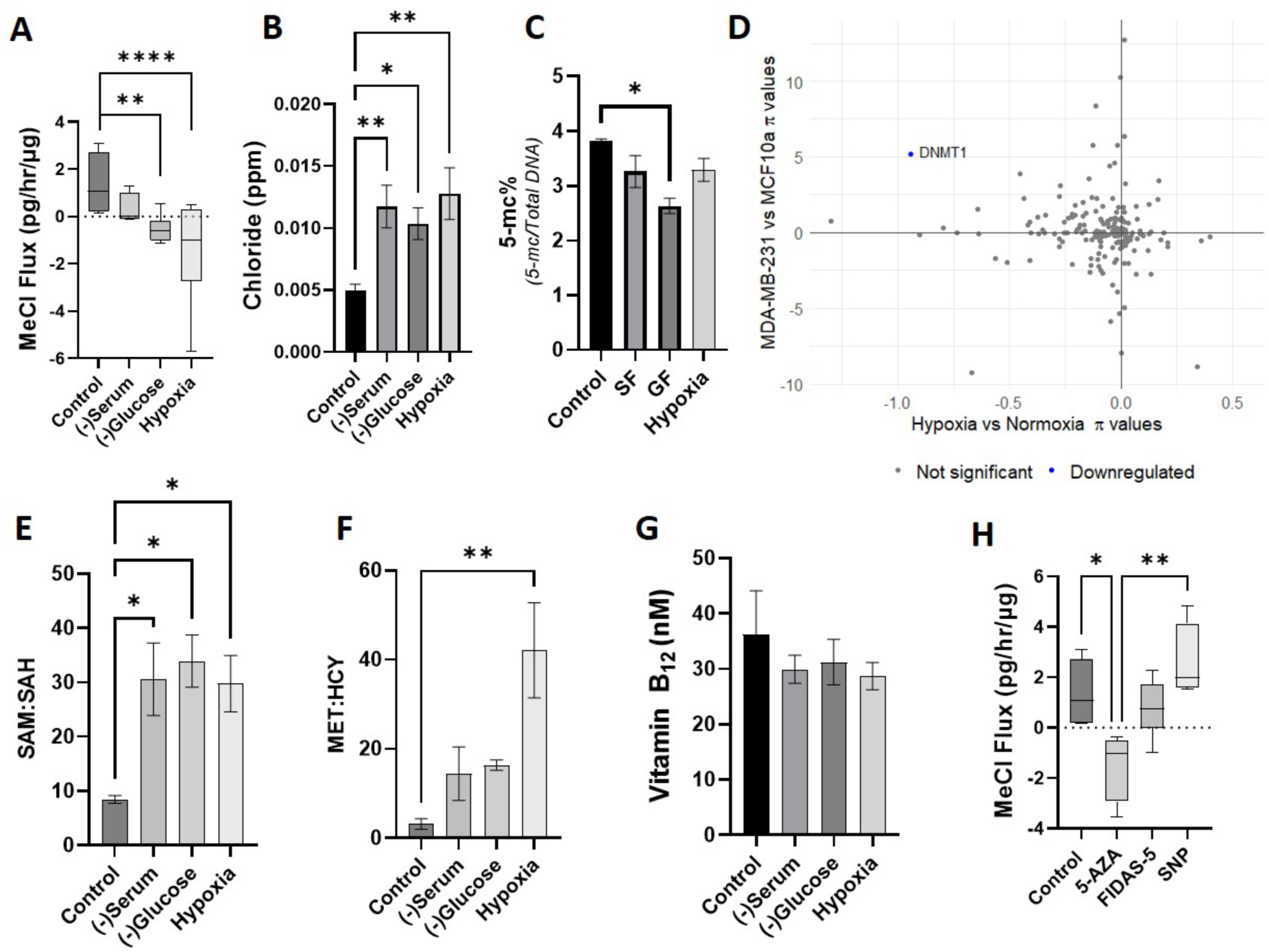
In MDA-Mb-231 cells methyl chloride (MeCl) fluxes correlate positively with changes in methylation metabolism and inversely with intracellular chloride content. **(A)** MeCl flux (pg MeCl/μg protein/hr) of MDA-MB-231 cells under plentiful resource conditions, without (-) serum/glucose, or under hypoxia. Data presented from figure 1 and hypoxia data from previously published data for comparative purposes, statistics for **A** are from Two-way ANOVA in the presence of the whole dataset (n=6). **(B)** Intracellular chloride content of cells under starvation/hypoxia conditions in parts per million (ppm; g Cl/g total protein) (n=4). **(C)** DNA methylation of cells under starvation/hypoxia conditions given as percentage 5-methylcytosine of total DNA (n=4). **(D)** Pi-chart of RNA data, comparing MDA-MB-231 (upwards on y-axis), MCF10a (down on y-axis), under normoxia (right, x-axis) and hypoxia (left, x-axis). **(E)** Ratio of S-adenosyl-methionine (SAM) to s-adenosyl-homocysteine (SAH) for cells under starvation conditions (n=4). **(F)** Ratio of methionine (MET) to homocysteine (HCY) for cells under starvation conditions (n=4). **(G)** Vitamin B12 (nM) for cells under starvation/hypoxia conditions (n=4). **(H)** MeCl flux (pg MeCl/μg protein/hr) of cells treated with 10 μM 5-azacytadine (5-AZA), 10 μM FIDAS-5 or 400 μM sodium nitroprusside (SNP) (n=6). Bar plots shown with mean ± SEM. One-way ANOVA with Tukey post hoc test was performed for **B, C, E-H**; * *p* < 0.05; ** *p* < 0.01; **** *p* < 0.0001.

Specifically, while in unstressed breast cells cellular chloride was correlated with MeCl flux (Figures 3A and B; suggesting a potential link between available chloride ions and methyl chloride production), in MDA-MB-231 cells under stress chloride intracellular content was inversely correlated with reduced production and/or consumption of MeCl (Figure 4B). In stressed MDA-MB-231 cells the difference in flux, from production to consumption, provides roughly the same amount of chloride ions over time as that increase observed in the cells, suggesting a metabolic utilization of methyl groups that leaves a chloride ion behind (e.g. −CH3Cl → CH3^+^ + Cl^−^).

All starvation/hypoxia conditions for MDA-MB-231 cells showed reduction in global DNA methylation potential, with significance found in the glucose starvation condition alone (Figure 4C), correlating with results seen in Figure 3, and suggesting that resource starvation is at least partially responsible for the observed reduction in methyl chloride fluxes.

Interestingly, differential analysis between MDA-MB-231 cells grown in hypoxic rather than normoxic conditions found only one significantly different gene: downregulation of *NOP2* (log2FC= −1.30; q=1.81) in hypoxia. Whilst no other genes met our significance thresholds (Figure S4A), including *DNMT1* (log2FC= −0.92; q=1.73), the data suggested an overall downregulation of methyl-transferase genes in hypoxia, supported by GSEA (NES: −1.701; FDR q: 0.0001, Figure S4B). Under normoxic conditions, MCF10a MeCl levels are lower than MDA-MB-231 (Figure 2). Thus, we investigated the similarities between MT gene expression in MCF10a with hypoxic MDA-MB-231s using a π-plot (Figure 4C). The only significantly changed gene was DNMT1, which was significantly down-regulated in both i) hypoxic conditions and ii) in MCF10a cells when compared with MDA-MB-231 in normoxic conditions.

Methyl transferase enzymes that produce MeCl utilize SAM (s-adenosylmethionine) as the methyl donor, with s-adenosylhomocysteine (SAH) as the by-product of the reaction. This allows for the ratio of SAM and SAH to represent methylation potential in cells [54, 55]. SAM:SAH ratio increased in all cell lines under starvation/hypoxia conditions (Figure 4E) suggesting either an underutilization of SAM during resource deprivation or an enhanced use of SAH. SAH is described as a potent inhibitor of methylation [54] and decreases relative to SAM show potential for methylation increasing and less consumption of SAM. Levels of SAM and SAH are shown in (Figure S4C, D).

### Consumption of MeCl correlates with metabolic potential of methionine

The observed dichotomous production/consumption of MeCl suggests two entirely different metabolisms in use by these cells. One metabolism is defined by methylation reactions that produce MeCl. The other, consumptive metabolism would use MeCl as a metabolic substrate rather than a metabolic output. As with methyl chloride production, metabolic utilization of methyl chloride has not been described or detailed in humans. Use of MeCl as a substrate has, however, been demonstrated and described in methylotrophic bacteria, via the *cmuA* methyltransferase [21, 45, 56]. The chloromethane methyltransferase protein (UNIPROT ID: F8J7J8 [45]) was therefore compared to the human proteasome via UNIPROT BLAST. The top result with 31.08% alignment was methionine synthase (MTR, UNIPROT ID: Q99707), alignment for these two proteins was then performed (Figure S5). High conservation was observed, with some conserved regions lying in the binding domains of the MTR structure, providing confidence that similar substrates, including methyl chloride, are used for both proteins.

MTR generates methionine from homocysteine, using vitamin b12 as a methyl donor. To investigate if MeCl consumption was related to methionine metabolism we investigated MTR, methionine, homocysteine and vitamin B12 behaviour of cells under starvation/hypoxia stress. Similar behaviour was seen for methionine (Figure S4E) and homocysteine (Figure S4F), as those observed for SAM and SAH. The ratio of methionine to homocysteine (MTH:HCY) increased for all starvation/hypoxia conditions with significance seen for cells in hypoxic environments (Figure 4F). In contrast, Vitamin B12 levels did not change significantly under starvation/hypoxia conditions (Figure 4G).

### Preventing methylation with 5-Azacytadine generates similar MeCl flux outcomes as starvation/hypoxia

Because results indicated a role for methylation metabolism in MeCl production, connected to alterations in SAM:SAH and MTH:HCY ratios, unstressed MDA-MB-231 cells were treated with 5-AZA (to block methylation [57]), FIDAS-5 (to block cellular generation of SAM [58]) and SNP (which has been shown to block methionine synthase [29]). 5-AZA treatment clearly generated a MeCl uptake response (Figure 4H), similar to effects of glucose or hypoxic starvation (Figure 2 and 3A). FIDAS-5 produced no change in MeCl flux and SNP treatment generated a slight, but insignificant increase in MeCl flux relative to unstressed MDA-MB-231 cells (Figure 4H). Viability of cells under these conditions was investigated using trypan blue assay. 5-AZA and SNP treatment produced a significant reduction in viability of 10-20% (Figure S4G).

### S-adenosylmethionine (SAM) treatment recovers methyl chloride flux by reversing methylation response and methionine/homocysteine behaviours under starvation

Because MeCl release by cells was suspected of being a product of methyl transferase activity, treatment with a primary substrate for methyl transferases, SAM, was performed for cells under starvation and MeCl fluxes quantified. In contrast to untreated cells, treatment of MDA-MB-231 (starved, hypoxic and control) with SAM resulted in no significant differences in MeCl fluxes between stressed and unstressed cells (Figure 5A). While there was an observable decrease in MeCl within all treatments compared to control, glucose starved cells which, when untreated, actively consumed MeCl, no longer consumed MeCl under SAM treatment (Figure 5A).

**Figure 5.**
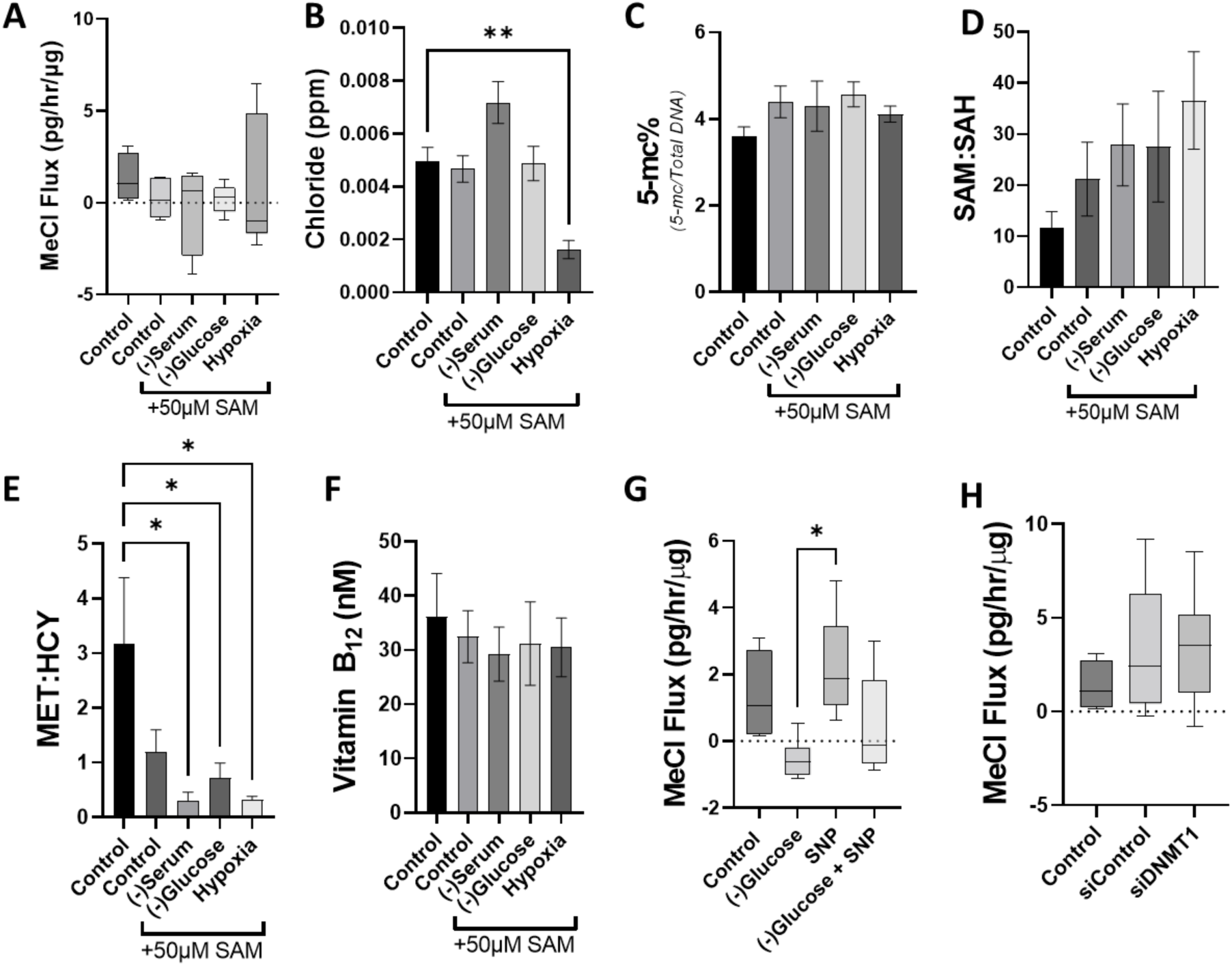
SAM treatment recovers methionine metabolic status and MeCl flux in starvation/hypoxia conditions. **(A)** MeCl flux (pg MeCl/μg protein/hr) of MDA-MB-231 cells treated with/without S-adensylmethionine (SAM) under control conditions or with S-adenosylmethionine and without (-) serum, glucose, oxygen. (n=6). **(B)** Intracellular chloride content of cells under starvation/hypoxia conditions or control, treated with SAM, in part per million (ppm; g Cl/g total protein) (n=4). **(C)** DNA methylation of cells under starvation/hypoxia conditions or control, treated with SAM, given as percentage 5-methylcytosine of total DNA (n=4). **(D)** Ratio of SAM to s-adenosyl-homocysteine (SAH) for cells under starvation/hypoxia conditions or control, treated with SAM (n=4). **(E)** Ratio of methionine (MET) to homocysteine (HCY) for cells under starvation/hypoxia conditions or control, treated with SAM (n=4). **(F)** Vitamin B12 (nM) for cells under starvation/hypoxia conditions or control, treated with SAM (n=4). **(G)** MeCl flux (pg/hr/μg) of glucose starved cells treated with 400 μM sodium nitroprusside (SNP) (n=6). **(H)** MeCl flux (pg/hr/μg) of untreated cells or those treated si-Control or si-DNMT1 (n=6). Bar plots shown with mean ± SEM. One-way ANOVA with Tukey post hoc test was performed. * *p* < 0.05; ** *p* < 0.01; *** *p* < 0.001; **** *p* < 0.0001.

Control and glucose starved cells show no change in chloride content following SAM treatment (Figure 5B). Serum starvation with SAM treatment produced a slight accumulation of chloride (Figure 5B), but to a lesser extent than serum starvation alone (Figure 4B). Cells in hypoxia with SAM supplementation, showed a significant reduction in chloride accumulation (Figure 5B), reversing the enhancements observed in untreated hypoxic cells (Figure 4B). SAM supplementation, in all cases, reduced the accumulation of chloride in resource stressed MDA-MB-231. SAM supplementation in serum and glucose starvation media produced no significant changes in media background volatile profiles (Figure S6A-C).

SAM is synthesised by methionine adenosyltransferase (MAT) [59]. Therefore, MAT levels were investigated in starvation and SAM supplemented MDA-MB-231 cells. Starvation of serum and glucose revealed a lower level of expression, shown by western blot (Figure S6D), which was significant when quantified (Figure S6E). SAM supplementation alone revealed little change compared to control while starvation with SAM revealed a similar trend, however slightly reduced (Figure S6D and E).

Cells under starvation showed no significant changes in DNA methylation when treated with SAM (Figure 5C). Methylation potential (SAM:SAH ratio) increased in cells treated with SAM and was further increased in cells under starvation/hypoxia (Figure 5D), maximum ratios of SAM:SAH when treated with SAM were similar to those seen in cells under starvation without SAM (Figure 4E). Intracellular SAM, increased to a lesser extent in cells under starvation with SAM supplementation (Figure S6H) than cells under starvation alone (Figure S4C). Effects upon intracellular SAH were also reduced in cells under starvation with SAM supplementation (Figure S6I) than those under starvation alone (Figure S4D).

Methionine to homocysteine ratios in MDA-MB-231 cells were significantly reduced under starvation/hypoxia conditions with SAM treatment (Figure 5E) in stark contrast to the increases seen in cells under resource stress alone, where this ratio increased (Figure 4F). Methionine levels increased, but not significantly in SAM treated cells (Figure S6J), as with cells under starvation alone. However, homocysteine levels substantively increased under SAM treatment (Figure S6K), whereas they decreased under resource stress alone (Figure S4F). Further to this, methionine synthase (MTR) levels from public RNA data, show no change under hypoxia compared to normoxia (Figure S7A). Vitamin B12 levels did not change under treatment with SAM (Figure 5F).

To determine if consumption of MeCl, which is pronounced in glucose starved cells, is a result of MTR activity, cells were treated with sodium nitroprusside (SNP) in control and glucose starvation conditions (Figure 5G). SNP treatment produced a slight increase in MeCl flux, a similar amount to that consumed under glucose starvation (Figure 5G and 4A). Treatment of glucose starved cells with SNP effectively removed consumption of MeCl by these cells (Figure 5G).

This data shows that SAM appears to be directly linked to MeCl production but is not an important component of methionine metabolism.

### Knockdown of DNMT1 does not alter MeCl flux in MDA-MB-231

To determine if DNMT1, the most abundant DNA methylating enzyme in humans [60] and most heavily researched methyl transferase in cancer [61–63], was the primary driving mechanism in MeCl production, Cells were treated with siRNA for DNMT1 or a si-scramble as control. Knockdown was successful, shown by western blot (Figure S7B), however no significant change was observed in MeCl flux under resource replete conditions (Figure 5H). This, along with observations in PRMT1, lead us to suggest that MeCl could be a product of multiple methyl transferase activities.

### Glutathione and reactive oxygen species are not linked to methyl chloride flux under starvation

Glutathione synthesis is derived from trans-sulfurated methionine and is linked to epigenetic changes and methylation activity [64, 65]. Glutathione levels were investigated to see if their levels corresponded with MeCl flux. In glucose starvation, glutathione reduced compared to serum and control conditions (Figure S7C). Treatment with SAM did not alter glutathione levels in control cells and produced no changes under serum or glucose starvation (Figure S7C). Therefore, alterations in MeCl fluxes were not well correlated with glutathione.

Reactive oxygen species (ROS) have been linked to VOCs in the breath and in cellular headspace [1]. Furthermore, while glutathione acts as a ROS scavenger, ROS may increase without changes in glutathione concentration and a main atmospheric sink for MeCl is radical attack [66]. To check for changes in ROS, as increases could be responsible for observed increases and decreases in VOCs, an amplex red assay was performed for cells under starvation conditions and with SAM treatment. Previously we have shown a significant reduction in ROS in MDA-MB-231 cells grown in hypoxic conditions for 24 h [9]. Under serum and glucose starvation following 24 h, ROS was slightly elevated compared to control (Figure S7D). Treatment with SAM produced a slight increase in ROS in control cells, a significant increase in serum starved and no change in glucose starvation (Figure S7D). Together with previously published results [9], these results suggest the VOCs studied here are not directly associated with variations in cellular ROS.

## Discussion

### Developing a conceptual model of methyl chloride metabolisms within cells

Cellular volatiles are significantly altered under serum, glucose and oxygen starvation. Consistently, methyl chloride (MeCl), which is produced by resource replete cells, is taken up under starvation and hypoxic stress and this uptake is matched by intracellular chloride accumulation. Variations in baseline (‘normal’) cell type MeCl release correlates with DNA methylation, but not activity of the ‘methyl-transferase-ome’ as a whole. Starvation and hypoxia induce MeCl uptake, with reduction in global DNA methylation, increases in methylation potential and changes in associated metabolites consistent with these outcomes. Publicly available RNA sequencing data supports these findings by revealing DNMT1 as a commonly reduced gene between starvation/hypoxia conditions and cell types where MeCl is reduced. Blocking methylation activity in cells produces a switch from production to uptake of MeCl but knockdown of DNMT1 produced no change. RNA data reveals significantly reduced enrichment of methyl transferases under starvation/hypoxia conditions, suggesting MeCl production is a result of global methyl transferase activity and not DNMT1 alone. This is further supported by supplementation with SAM in starvation conditions lessening the effect of starvation upon MeCl fluxes. Further to this, active consumption of MeCl is linked to methionine synthase, which is supported by methionine synthase blocking agents removing starvation effects upon MeCl fluxes.

Volatile flux linked to cellular mechanistic processes has been shown in this research for MeCl, both as a product of methyl transferase activity and as a substrate for methionine synthesis. To highlight these disparate processes, a schematic is presented in Figure 6. Here, the activity of methyl transferases generates MeCl while consumption of MeCl is driven by methionine synthase, replacing vitamin b12 as a methyl donor, resulting in the observed accumulation of chloride ions. We have shown that MeCl levels rise and fall alongside DNA methylation levels, however, knockdown of DNMT1, the primary enzyme for DNA methylation, did not significantly alter MeCl flux. Protein alignment of MeCl producing enzymes in plants with the human database revealed alternate candidate SAM-dependent methyl transferases, including TPMT. We determined that DNA methylating enzymes alone may not drive MeCl production as publicly available data shows a down regulation of methyl transferases under hypoxic conditions and treatment with 5-AZA, which blocks a range of methylation events, prevented release of MeCl. Furthermore, treatment with SAM prevented loss of MeCl production under starvation conditions. Additional SAM, in this instance, may promote methyl transferase activity, increasing MeCl production. This is supported by published evidence that cells under hypoxic conditions are hypo methylated [67], potentially through the impairment of the production of SAM in hypoxic conditions [68, 69].

**Figure 6.**
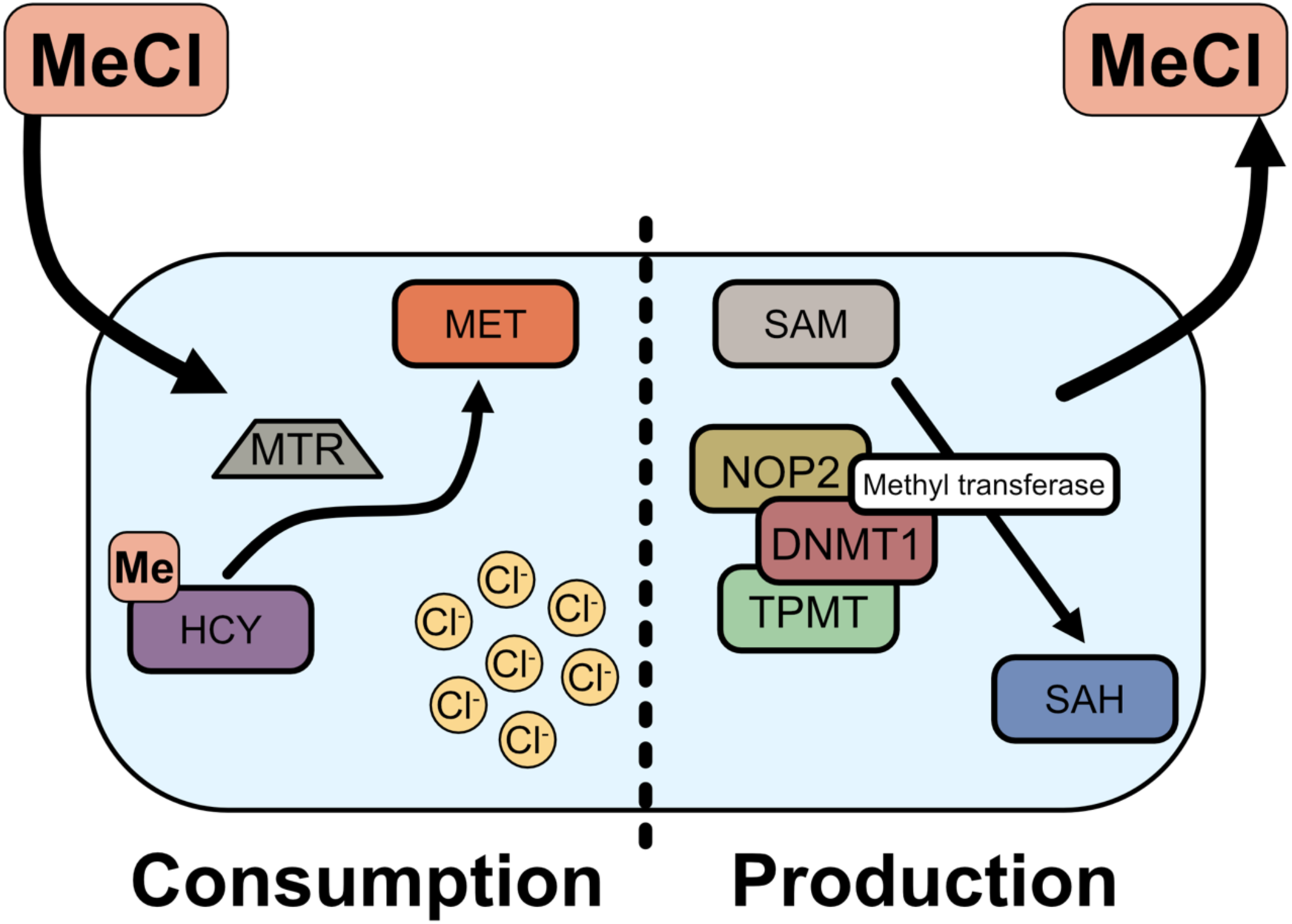
Proposed mechanisms of methyl chloride (MeCl) consumption and production. MeCl consumption by cells is through the action of methionine synthase (MTR), generating methionine (MET) through the addition of a methyl group from MeCl to homocysteine (HCY), resulting in an accumulation of chloride ions. Activity of S-adenosylmethionine (SAM) dependent methyl transferases which use SAM as a methyl donor, producing S-adenosyl-L-homocysteine as a primary, and MeCl as a secondary, metabolic product.

### Implications for diagnosis and treatment of breast cancer

To discover biomarkers capable of distinguishing cancer in the breath of tumour bearing mice and ultimately in the breath of breast cancer patients this research has utilised a methodology capable of measuring cellular metabolism of VOCs in models mimicking the tumour pathological environment, namely serum, glucose and oxygen starvation. The primary compound of interest arising from theses studies, methyl chloride, is established here as an important identifier of the SAM, SAH, methionine and homocysteine status of the cell. This provides an important overview of the methylation potential of the system under study.

DNA methylation profiles within cancer, manifesting as hypomethylation across the genome and hypermethylation for specific loci, present a current emerging technology for cancer diagnostics [62, 70], with large scale ongoing human trials [63, 71]. We present, through this work, methyl chloride (MeCl) flux - a direct indicator of cellular methylation status and related metabolic pathways - as novel biomarkers represented in human breath

MeCl, is a known methylating agent and is exhaled in human breath in the range of 2.5 to 33 parts per billion by volume (ppbv), up to ∼60 times that of inhaled air [16]. Isoprene is the most studied VOCs in human breath [72], with reported concentrations of 100ppbv (at rest) and much higher concentrations following exercise [72]. Compared to isoprene and other compounds such as acetone (1.2-1880 ppbv), methanol (160-2000 ppbv) and ethanol (13-1000 ppbv) [73], MeCl, is found in lower concentrations (3.5 ± 2.4 ppbv) but is still a substantive component of human breath [16, 17, 74]. Furthermore, while our pilot study results for MeCl did not achieve statistical significance at the 0.05 level, the observed trends (p<0.09) warrant further investigation.

Because starvation of glucose, hypoxia and treatment with 5-AZA not only prevented production of MeCl but generated an active consumption by cells, we propose that MeCl is both used and produced by cells. This “pull” component of our ‘push-pull’ model presents a novel method for methionine generation, an essential amino acid for protein synthesis and other biochemical reactions required for cell viability and growth [65]. Methionine is used heavily by cancer cells in growth and response to stress [65] and the stress responses shown here may be linked to this dependence. MDA-MB-231 cells under hypoxic conditions have been shown to have increased levels of methionine [75], in line with our findings. Furthermore, we have shown cells under various stresses consuming MeCl, including chemotherapeutic stress with Doxorubicin [5], which has been shown to increase intra and extra cellular methionine in skeletal muscle cells [76].

A balance between production and consumption of MeCl is observed in the breath of mice in this study. Mice with larger tumours (e.g.- with increased levels of cells persisting in conditions with low nutrient delivery, or resource stress) have reduced levels of MeCl compared to smaller tumours, suggesting that greater resource stress within the tumours leads to a more complete transition from one metabolic state to the other (Figure 1E). This is further supported by the observation that in the breath of breast cancer patients we see this same trend coupled with a significant increase in chloroform, which we propose arises from the chronic accumulation of intracellular chloride driven by sustained MeCl consumption [18]. The duration of cancer-associated cellular stress may therefore be a critical determinant of the chloroform signal: short-term (24 h) cell line starvation produces chloride accumulation without measurable chloroform release, intermediate-duration (weeks) tumour growth in mice produces a modest chloroform increase, and long-standing (months to years) human cancers produce a significant chloroform signature. This time-dependence is consistent with chloroform formation being a downstream consequence of the chloride accumulation predicted by our push-pull model (Figure 6), reaching detectable equilibrium only after sustained metabolic perturbation.

Increased methionine levels and decreased methylation in cells under the stress of the tumour microenvironment [65, 67], may explain reduced MeCl in the breath of tumour bearing mice. MeCl consumption through methionine synthesis coupled with a decrease in release through reduced methylation may explain the observed trends, supporting the mechanism presented in figure 6. Furthermore, intra-cellular methionine levels have been shown to increase with tumour size (Kawaguchi et al. 2018).

## Conclusion

The methodology described here for both mouse breath (static headspace) cellular headspace, and human patient breath as previously described [5, 9, 18], takes two headspace samples to determine flux of VOC over time. This methodology is substantively different from the majority of breath and cellular VOC approaches which often concentrate VOCs onto materials for thermal desorption of a single time point, providing a concentration estimate [1]. While the presented method is less automated, it allows description of active consumption or production of VOCs, which we have shown here to be important, descriptive characteristics of stress. In mice breath for example, while levels of butanone in the breath of tumour bearing mice is not significantly different from zero, it is significantly different from control, where mice are consuming this VOC.

Through modelling tumour environment in cells, measuring select VOCs and then testing the breath of tumour bearing mice and breast cancer patients for those same targets, we have identified two VOCs (MeCl and butanone) out of seven initial targets which were significantly altered in the cellular model (Figure 2) and which translated to the mouse model (Figure 1) and ultimately to the human (Figure 1 and [18]). Chloroform emerges as a third translational biomarker whose signal strengthens with the duration of cancer-associated stress, becoming most pronounced in the breath of human patients. We have then described how observed changes in MeCl flux (Figure 6) can be explained within the context of methylation and methionine synthesis, both of which are altered in cancer [61, 65, 77].

## Author Contributions

Conceptualization, T.I., W.J.B. and K.R.R.; Data curation, T.I.; Formal analysis, T.I., A.M.; Funding acquisition, T.I., J.T., J.P., S.T.S., W.J.B. and K.R.R.; Investigation, T.I.; Methodology, T.I., A.M. and K.R.R.; Project administration, T.I. and S.T.S.; Resources, W.J.B.; Visualization, T.I., A.M.; Writing—original draft, T.I.; Writing—review and editing, S.T.S., W.J.B., M.R. and K.R.R.

## Acknowledgements

The authors would like to acknowledge the support provided by Mark Bentley in the University of York Department of Biology workshop and the University of York Technology Facility. The authors would also like to recognise the ongoing core support to WB from York Against Cancer.

## Ethical Statement

Human breath studies received ethical approval from an NHS Research Ethics Committee (IRAS ID: 318636) and University of York Biology Ethics Committee (reference: KR202302). All participants provided written informed consent before any study procedures were undertaken. The study adhered to the ethical principles outlined in the Declaration of Helsinki and Good Clinical Practice guidelines. Approval covered patient recruitment, breath collection, data handling, and analysis as described in the protocol.

