## Supplementary for "Breast cancer cohort study identifies an effective, non-invasive, breath-based diagnostic linked to cellular environment-dependent, novel methylation metabolisms"

Supplementary Material

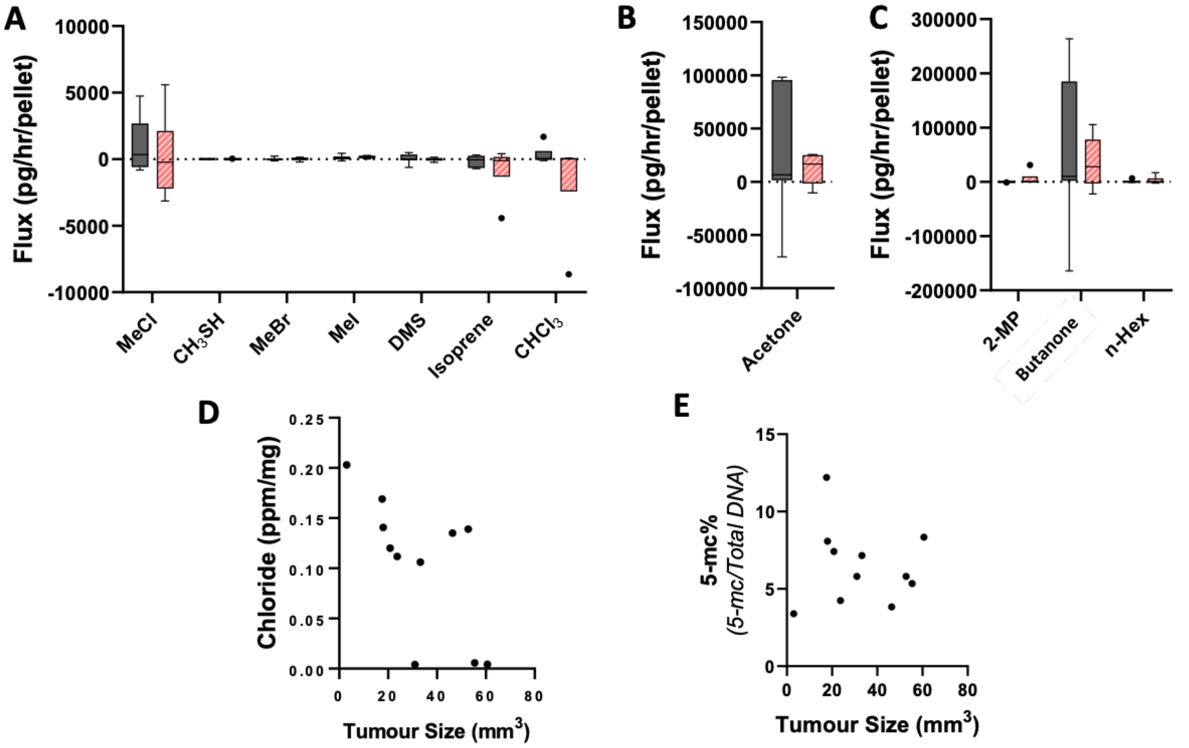


Supplementary Figure 1**. Volatile flux and tumour chloride and DNA methylation content from MDA-MB-231 tumour xenograft bearing mice. (A-C)** . Volatile flux (g/hr/pellet) of mouse faecal pellets from control and MDA-MB-231 xenograft tumour bearing mice. Normalised to number of pellets (n=6). **(D)** Chloride content in parts per million (ppm) normalised to weight of sample in mg compared to total tumour size in $mm^3$. **(E)** 5-methyl cytosine (5-mc) content as percentage of total DNA for tumours compared to total tumour size in $mm^3$.

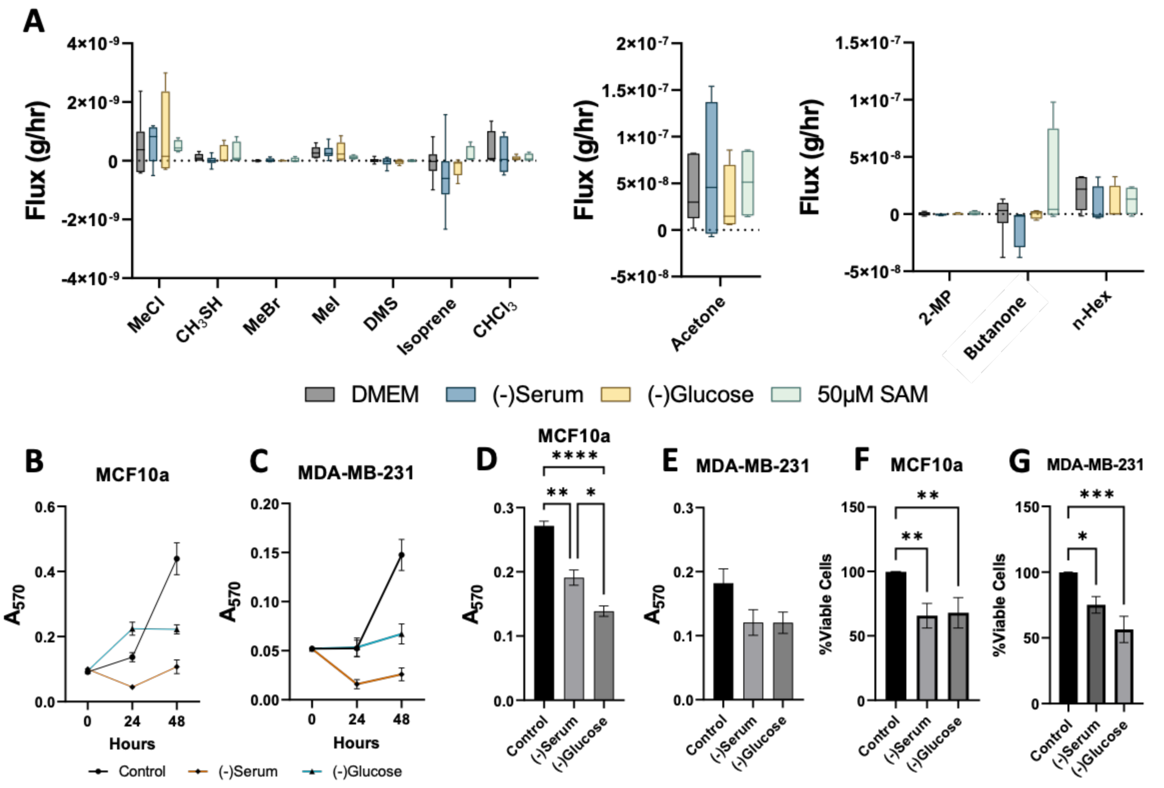


Supplementary Figure 2**. Volatile flux for media backgrounds and cell response to starvation.** Select volatile compound flux (g/hr[-1]) for media control conditions **(A).** SAM, S-adensolymethionine; $CHCl_3$, chloroform; DMS, dimethyl sulfide; MeBr, methyl bromide; MeCl, methyl chloride; MeI, methyl iodide; MeSH, methanoethiol; 2-MP, 2 methyl pentane; n-Hex, n-hexane. Boxplot whiskers show median ± Tukey distribution (n=6). **(B,C)** Cell growth curves measured by sulphorhodamine B assay for MCF10a **(B)** and MDA-MB-231 **(C)** (n=3). **(D)** MTT assay for cells following 24 hour starvation (n=3). **(D)** Cell viability/death measured by trypan blue assay for MCF10a **(F)** and MDA-MB-231 **(G)** (n=3). Mean One-way ANOVA with tukey post hoc analysis performed for **D-G,** error bars are mean ± SEM, *$p < 0.05$; **$p < 0.01$; ***$p < 0.001$; ****$p < 0.0001$.

**A**

```
tr|Q9ZSZ7|Q9ZSZ7_BATMA      MSTVANIAPVFTGDCKTIPTP--EECATFLYKVVNSGGWEKCWVEEVIPWDLGVPTPLVL    58
sp|P51580|TPMT_HUMAN        ----------MDGTRTSLDIEEYSDTEVQKNQVLTLEEWQDKWVNGKTAFHQEQGHQLLK    50
                                      : *  .::     .:  .   :*:.   *:. **:    :.      *:

tr|Q9ZSZ7|Q9ZSZ7_BATMA      HLVKN--NALPNGKGLVPGCGGGYDVVAMANPERFMVGLDISENALKK---------ARE    107
sp|P51580|TPMT_HUMAN        KHLDTFLKGKSGLRVFFPLCGKAVEMKWFADRGHSVVGVEISELGIQEFFTEQNLSYSEE    110
                            : :..  :.  . : :.* ** . ::  :*:  : :**::*** .:::         :.*

tr|Q9ZSZ7|Q9ZSZ7_BATMA      TFSTMPNSSCFSFVKE-------DVF--TWRPEQPFDFIFDYVFFCAIDPKMRPAWGKAM    158
sp|P51580|TPMT_HUMAN        PITEIPGTKVFKSSSGNISLYCCSIFDLPRTNIGKFDMIWDRGALVAINPGDRKCYADTM    170
                             :: :*.:. *.  .        .:*         **:*:*   : **:*  *  .:..:*

tr|Q9ZSZ7|Q9ZSZ7_BATMA      YELLKPDGELI--TLMYPITNHEGGPPFSVSESEYEKVLVPLGFKQLSLEDYSDLAVEPR    216
sp|P51580|TPMT_HUMAN        FSLLGKKFQYLLCVLSYDP-TKHPGPPFYVPHAEIERLFGKICNIR-CLEKVDA------    222
                            :.**  . : :  .* *   .:. **** * .:* *:::  :   : .**. .

tr|Q9ZSZ7|Q9ZSZ7_BATMA      KGKEKLARWKKMNN---------- 230
sp|P51580|TPMT_HUMAN        -FEERHKSWGIDCLFEKLYLLTEK 245
                              :*:   *
```

**B**

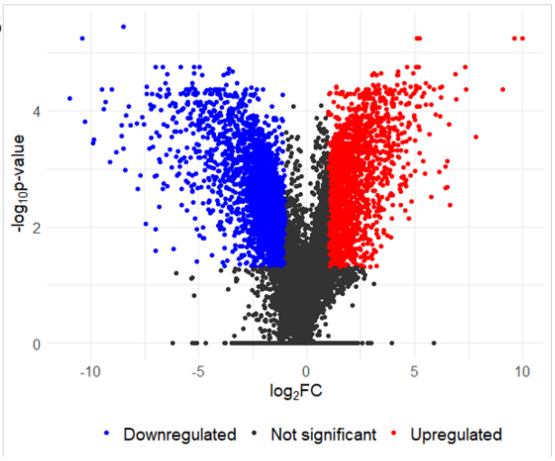


Supplementary Figure 3**. Protein alignment for TPMT and methyl chloride transferase and volcano plot. (A)** Alignment results from UNIPROT for human TPMT (thiopurine methyltransferase, UNIPROT ID: P51580) and plant methyl chloride transferase (*Batis maritima,* UNIPROT ID: Q9ZSZ7*).* Alignment results revealed 24.8% similiarity, the most similar protein in the UNIPROT human proteasome against methyl chloride transferase. ***** = same residue **:** , **.** =closely linked residues. Underlined and red sections show binding domains. **(B)** Volcano plot of RNA seq data for -log10 LRT (likelihood ratio test) q values (corrected p values using Benjamini-Hochberg) vs mean log2 fold change of MDA-MB-231 vs MCF10a.

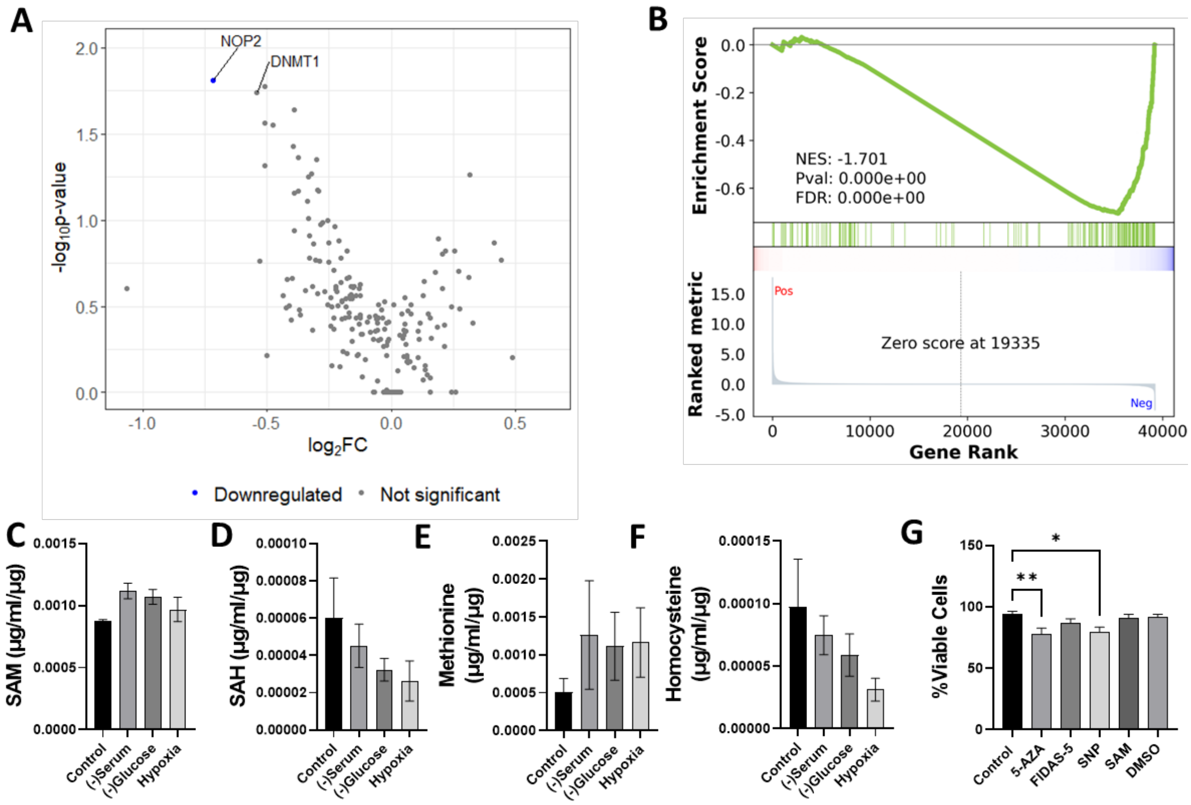


Supplementary Figure 4**. Effects of starvation upon mRNA and methylation metabolites in MDA-MB-231. (A)** Volcano plot of RNA seq data for -log10 LRT (likelihood ratio test) q values (corrected p values using Benjamini-Hochberg) vs mean log2 fold change of hypoxia vs normoxia. **(B)** Gene set enrichment analysis results of RNA data from **(A)**. S-adenosylmethioine (SAM) **(C)**, S-adenosyl-L-homocysteine (SAH) **(D)**, Methioine **(E)** and homocysteine **(F)** intracellular MDA-MB-231 content in µg/ml/ug, for cells under starvation conditions or control. **(G)** Percentage viable cells for MDA-MB-231 cells following 24 hours treatment with 10 µM 5-azacytadine (5-AZA), 10 µM FIDAS-5 or 400 µM sodium nitroprusside (SNP), 50 µM SAM or 0.00005% DMSO measured by Trypan blue assay. Bar plots shown with mean ± SEM, n=4 **.** One-way ANOVA with Tukey post hoc analysis performed for all bar charts. *$p < 0.05$; **$p < 0.01$.

```
tr|F8J7J8|F8J7J8_HYPSM          -------------MTQVPKMTSRERLFA--------------------------------    15
sp|Q99707|METH_HUMAN            KGLLDGGVDILLIETIFDTANAKAALFALQNLFEEKYAPRPIFISGTIVDKSGRTLSGQT    240
                                              * . . .::  ***

tr|F8J7J8|F8J7J8_HYPSM          -------------------------------AVTMQTLPDQVPCVP--------------    30
sp|Q99707|METH_HUMAN            GEGFVISVSHGEPLCIGLNCALGAAEMRPFIEIIGKCTTAYVLCYPNAGLPNTFGDYDET    300
                                                                :  :     * * *

tr|F8J7J8|F8J7J8_HYPSM          -LLMTRGIREG---GI----------TVDQ--ALRDGEASAHAKIKALKKFGGDVIIPGT    74
sp|Q99707|METH_HUMAN            PSMMAKHLKDFAMDGLVNIVGGCCGSTPDHIREIAEAVKNCKPRVPPATAFEGHMLLSGL    360
                                  :*:: :::    *:          * *:   : :.  ..: ::   . * *.::: *

tr|F8J7J8|F8J7J8_HYPSM          DLFTPVECVEGCELDYLPYAQPSLVKHPTPTKEAFYRYKEKYLREGFKPSERVLQIQKEA    134
sp|Q99707|METH_HUMAN            EPFRIG-----------PYTNFVNIGERCN-VAGSRKFAKLIMAGNYEEALCVAKVQVEM    408
                                : *              **::   : .      .  :: :  :  .:: :  * ::* *

tr|F8J7J8|F8J7J8_HYPSM          RTMIAQGVKDTHAMPTPVGGPITTAQLMTGSSEFLSYISDDPDYAKEVTELALDIVKNVC    194
sp|Q99707|METH_HUMAN            GA----QVLDVNMDDGMLDGPS-------AMTRFCNLIASEPDIAKVPLCI-----DSSN    452
                                 :     * *.:     :.**        . :.* . *:.:** **    :     ..

tr|F8J7J8|F8J7J8_HYPSM          RMMFEAGIDVCNILDPFNSSDILP--------PDTYREFGLPYQKRL-------------    233
sp|Q99707|METH_HUMAN            FAVIEAGLKCCQGKCIVNSISLKEGEDDFLEKARKIKKYGAAMVVMAFDEEGQATETDTK    512
                                  ::***:. *:    .** .:            . :::*

tr|F8J7J8|F8J7J8_HYPSM          -----FAYIKEIGGIGFTHTCTFTQPIWRDIANNGCFNFNGDMYP-GMDHAKRAIGGQ--    285
sp|Q99707|METH_HUMAN            IRVCTRAYHLLVKKLGFNPNDIIFDPNILTIGT-GM--EEHNLYAINFIHATKVIKETLP    569
                                      **   :  :**. .  : :*    *.. *    : ::*  .: **.:.*

tr|F8J7J8|F8J7J8_HYPSM          -ISLMGTLSPFSTLMHGSTTD---VANEVKKLAAEVGYNGGLIVMPGCDIDWTIPDENLK    341
sp|Q99707|METH_HUMAN            GARISGGLSNLSFSFRGMEAIREAMHGVFLYHAIKSGMDMGIVNA----GNLPVYDDIHK    625
                                   : * ** :*  ::*  :    : . .   * : * : *::       : : *:   *

tr|F8J7J8|F8J7J8_HYPSM          AMIETCASIKYPMDVAALGDLSNVYLAGHPKHPGKRAPSTAGDTDVAEAKTHHKELTPQQ    401
sp|Q99707|METH_HUMAN            ELLQLCEDLIWNKDPEATEKLLRYA--QTQ--------GT-GGKKVI----------QTD    664
                                 ::: * .: :  *  *  .* .               .* *...*             :

tr|F8J7J8|F8J7J8_HYPSM          EVNEKLVEAIMEYDGDKAIEW--------VKKGLERGMTAQDIVFDGLSLGMKVVGDMYE    453
sp|Q99707|METH_HUMAN            EWRNGPVEERLEYALVKGIEKHIIEDTEEARLNQKKYPRPLNIIEGPLMNGMKIVGDLFG    724
                                * .:  **  :**   *.**         .: . ::     :*: . *  ***:***::

tr|F8J7J8|F8J7J8_HYPSM          RNERFVTDMLKAAKTMDKAMPILTPLLEQAGG-----------DGGPTGTVVVGLVRGNT    502
sp|Q99707|METH_HUMAN            AGKMFLPQVIKSARVMKKAVGHLIPFMEKEREETRVLNGTVEEEDPYQGTIVLATVKGDV    784
                                 .: *: :::*:*:.*.**:  * *::*:              :.   **:*:. *:*:.

tr|F8J7J8|F8J7J8_HYPSM          QDIGKNLVCLMLKANGFKVIDLGKNVKPEQFIESAEKENAVAIGMSVMTNSST---VYVE    559
sp|Q99707|METH_HUMAN            HDIGKNIVGVVLGCNNFRVIDLGVMTPCDKILKAALDHKADIIGLSGLITPSLDEMIFVA    844
                                :*****:* ::* .*.*:*****  .  ::::::* ..:*  **:* : . *    ::*

tr|F8J7J8|F8J7J8_HYPSM          KVKELLDKAGKGDKYLLMCGGAAANK-GVADKMGVKYGLD------ANAAVSLVK----D    608
sp|Q99707|METH_HUMAN            KEMERL-----AIRIPLLIGGATTSKTHTAVKIAPRYSAPVIHVLDASKSVVVCSQLLDE    899
                                *  * *     . :  *: ***::.*  .* *:. :*.        *. :* : .    :

tr|F8J7J8|F8J7J8_HYPSM          HLQAAA------------------------------------------------------    614
sp|Q99707|METH_HUMAN            NLKDEYFEEIMEEYEDIRQDHYESLKERRYLPLSQARKSGFQMDWLSEPHPVKPTFIGTQ    959
                                :*:
```

Supplementary Figure 5**. Protein alignment of the methylotroph cmuA protein and human methionine synthase.** Alignment results from UNIPROT for human MTR (methionine synthase, UNIPROT ID: Q99707) and bacterial chloromethane (methyl chloride) methyltransferase (UNIPROT ID: F8J7J8*).* Alignment results revealed 31.08% similarity, the most similar protein in the UNIPROT human proteasome against chloromethane methyl transferase. *= same residue : , . =closely linked residues. Underlined and red sections show binding domains.

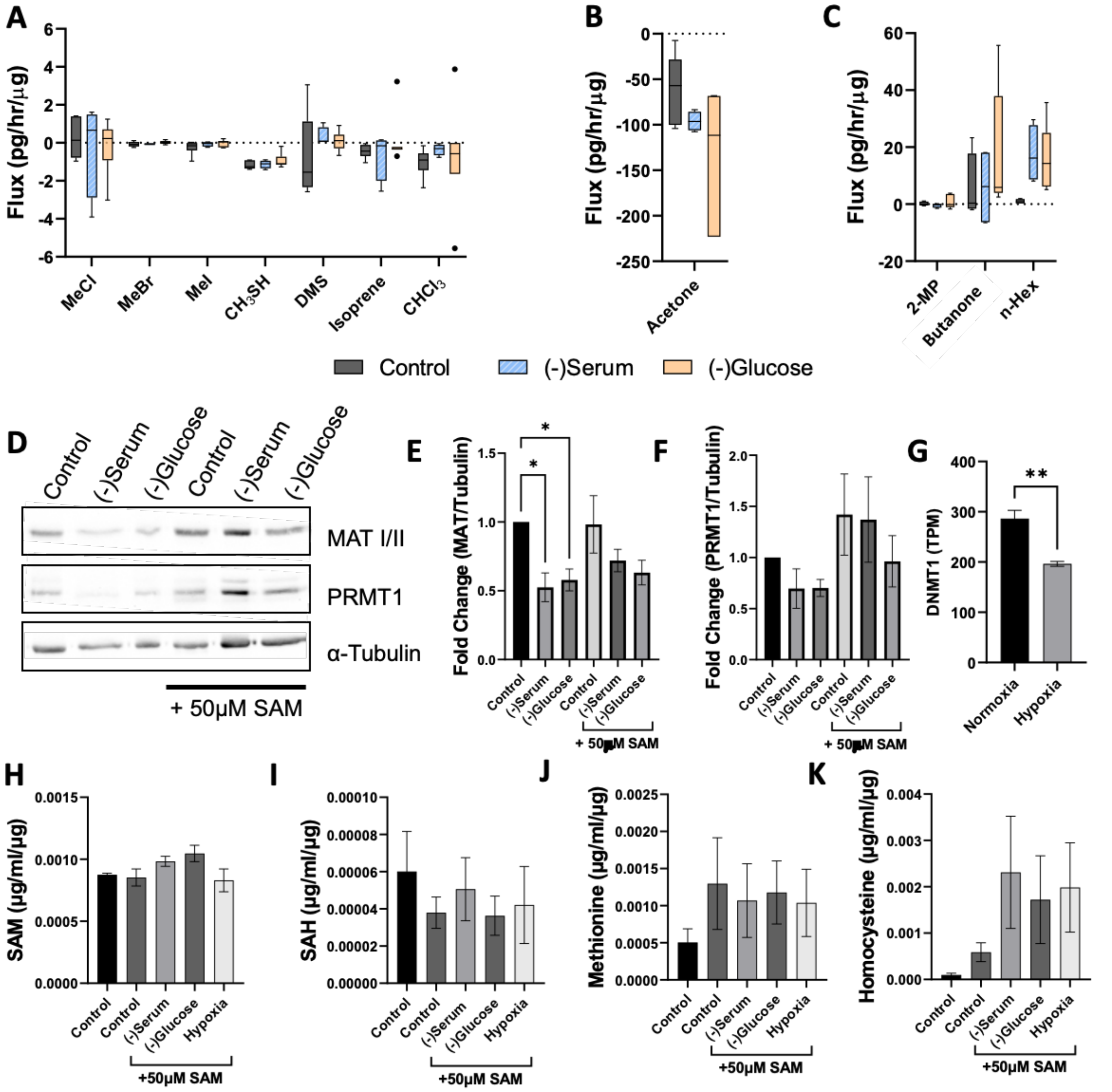


Supplementary Figure 6**. Effects of S-adenosylmethionine (SAM) treatment upon MDA-MB-231 cells. (A-C)** Volatile flux (pg/hr/µg) for MDA-MB-231 in control media, media without serum or media without glucose, all supplemented with 50 µM SAM. Media subtracted and protein-normalised. **(D)** Representative western blot of MDA-MB-231 cell lysates probed for MATI/II, PRMT1 and α-Tubulin in starvation conditions with or without SAM. **(E)** Quantification of MATI/II western blots by densitometry analysis of conditions in **D** normalised to α-tubulin and expressed as fold change compared to control **(F) )** Quantification of PRMT1 western blots by densitometry analysis of conditions in **D** normalised to α-tubulin and expressed as fold change compared to control. **(G)** DNMT1 RNA levels in transcripts per million (TPM) of MDA-MB-231 cells in normoxic or hypoxic conditions from publicly available data. S-adenosylmethioine (SAM) **(H)**, S-adenosyl-L-homocysteine (SAH) **(I)**, Methioine **(J)** and homocysteine **(K)** intracellular MDA-MB-231 content in µg/ml/ug, for cells under starvation conditions or control supplemented with SAM. (-)Serum = media without serum, (-)Glucose = media without glucose. $CHCl_3$ = Chloroform, DMS = Dimethyl sulphide, MeBr = Methyl bromide, MeCl = Methyl chloride, MeI = Methyl iodide, MeSH = Methanoethiol. Boxplot whiskers show median ± Tukey distribution (n=6). Two-way ANOVA with Bonferroni post hoc test was performed for **A-C**. Bar plots shown with mean ± SEM. One way ANOVA with Tukey posthoc analysis performed for bar charts **E, F, H-K**. Students T-test performed for **G**. *$p$ < 0.05; ***$p$ < 0.001.

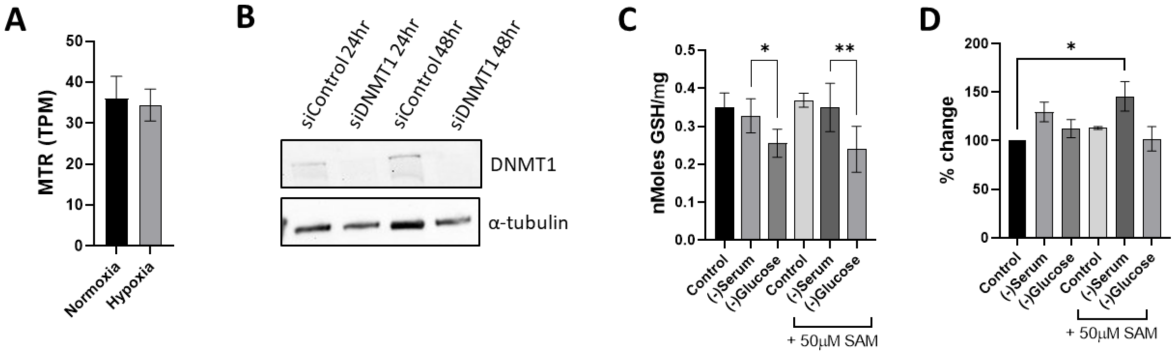


Supplementary Figure 7**. Glutathione and reactive oxygen species investigation and DNMT1 knockdown in MDA-MB-231. (A)** MTR (methionine synthase) RNA levels in transcripts per million (TPM) of MDA-MB-231 cells in normoxic or hypoxic conditions from publicly available data. (n=3) . **(B)** Representative western blot of MDA-MB-231 cell lysates probed for DNMT1 and α-Tubulin in treated with si-RNA targeting DNMT1 or scrambled control at 24 hours or 48 hours post treatment. **(C)** Glutathione content of MDA-MB-231 cells, normalised to protein content (GSH/mg) in nMoles, for cells in control, serum or glucose starvation with our with out S-adenslymethioine (SAM) treatment (n=3). **(D)** Amplex red assay for reactive oxygen species. Treatment conditions as with **C**, expressed as percentage change compared to control (n=3). Bar plots shown with mean ± SEM. One way ANOVA with Tukey posthoc analysis performed for bar charts **C** and **D**. *$p$ < 0.05; **$p$ < 0.01.

| ENSEMBL ID | Gene ID | Approved symbol | Approved name | Location |
|---|---|---|---|---|
| ENSG00000188573.8 | FBLL1 | FBLL1 | fibrillarin like 1 | 5q34 |
| ENSG00000169519.21 | METT5D1 | METTL15 | methyltransferase like 15 | 11p14.1 |
| ENSG00000244026 | FAM86D | FAM86DP | family with sequence similarity 86 member D, pseudogene | 3p12.3 |
| ENSG00000214756.8 | METTL12 | CSKMT | citrate synthase lysine methyltransferase | 11q12.3 |
| ENSG00000227835 | CARM1L | CARM1P1 | coactivator associated arginine methyltransferase 1 pseudogene 1 | 9p24.2 |
| ENSG00000107614.22 | TRDMT1 | TRDMT1 | tRNA aspartic acid methyltransferase 1 | 10p13 |
| ENSG00000100462.16 | PRMT5 | PRMT5 | protein arginine methyltransferase 5 | 14q11.2 |
| ENSG00000101654.18 | RNMT | RNMT | RNA guanine-7 methyltransferase | 18p11.21 |
| ENSG00000132275.11 | RRP8 | RRP8 | ribosomal RNA processing 8 | 11p15.4 |
| ENSG00000071462.12 | WBSCR22 | BUD23 | BUD23 rRNA methyltransferase and ribosome maturation factor | 7q11.23 |
| ENSG00000168806.8 | LCMT2 | LCMT2 | leucine carboxyl methyltransferase 2 | 15q15.3 |
| ENSG00000145194 | ECE2 | ECE2 | endothelin converting enzyme 2 | 3q27.1 |
| ENSG00000185238.13 | PRMT3 | PRMT3 | protein arginine methyltransferase 3 | 11p15.1 |
| ENSG00000241644.2 | INMT | INMT | indolethylamine N-methyltransferase | 7p14.3 |
| ENSG00000171806.12 | C1orf156 | METTL18 | methyltransferase like 18 | 1q24.2 |
| ENSG00000169093.16 | ASMTL | ASMTL | acetylserotonin O-methyltransferase like | Xp22.3_Yp11.3 |
| ENSG00000145002.13 | FAM86B2 | FAM86B2 | family with sequence similarity 86 member B2 | 8p23.1 |
| ENSG00000174912 | METT5D2 | METTL15P1 | methyltransferase like 15 pseudogene 1 | 3q25.31 |
| ENSG00000141744.4 | PNMT | PNMT | phenylethanolamine N-methyltransferase | 17q12 |
| ENSG00000093010.15 | COMT | COMT | catechol-O-methyltransferase | 22q11.21 |
| ENSG00000120265.19 | PCMT1 | PCMT1 | protein-L-isoaspartate (D-aspartate) O-methyltransferase | 6q25.1 |
| ENSG00000105202.9 | FBL | FBL | fibrillarin | 19q13.2 |
| ENSG00000130816.17 | DNMT1 | DNMT1 | DNA methyltransferase 1 | 19p13.2 |
| ENSG00000166741.8 | NNMT | NNMT | nicotinamide N-methyltransferase | 11q23.2 |
| ENSG00000111641.12 | NOP2 | NOP2 | NOP2 nucleolar protein | 12p13.31 |
| ENSG00000196433.13 | ASMT | ASMT | acetylserotonin O-methyltransferase | Xp22.3_Yp11.3 |
| ENSG00000169710.9 | FASN | FASN | fatty acid synthase | 17q25.3 |

| | | | | |
|---|---|---|---|---|
| ENSG00000150540.14 | HNMT | HNMT | histamine N-methyltransferase | 2q22.1 |
| ENSG00000160310.19 | PRMT2 | PRMT2 | protein arginine methyltransferase 2 | 21q22.3 |
| ENSG00000037474.15 | NSUN2 | NSUN2 | NOP2/Sun RNA methyltransferase 2 | 5p15.31 |
| ENSG00000130005.13 | GAMT | GAMT | guanidinoacetate N-methyltransferase | 19p13.3 |
| ENSG00000124713.6 | GNMT | GNMT | glycine N-methyltransferase | 6p21.1 |
| ENSG00000164603.12 | C7orf60 | BMT2 | base methyltransferase of 25S rRNA 2 homolog | 7q31.1 |
| ENSG00000126814.7 | TRMT5 | TRMT5 | tRNA methyltransferase 5 | 14q23.1 |
| ENSG00000291151 | NSUN5P1 | NSUN5P1 | NSUN5 pseudogene 1 | 7q11.23 |
| ENSG00000110871.15 | COQ5 | COQ5 | coenzyme Q5, methyltransferase | 12q24.31 |
| ENSG00000203791.15 | METTL10 | EEF1AKMT2 | EEF1A lysine methyltransferase 2 | 10q26.13 |
| ENSG00000162639.16 | C1orf59 | HENMT1 | HEN methyltransferase 1 | 1p13.3 |
| ENSG00000101247.18 | C20orf7 | NDUFAF5 | NADH:ubiquinone oxidoreductase complex assembly factor 5 | 20p12.1 |
| ENSG00000203740.4 | METTL11B | NTMT2 | N-terminal Xaa-Pro-Lys N-methyltransferase 2 | 1q24.2 |
| ENSG00000139780.8 | C13orf39 | METTL21C | methyltransferase 21C, AARS1 lysine | 13q33.1 |
| ENSG00000005194.15 | NSUN5C | NSUN5P2 | NSUN5 pseudogene 2 | 7q11.23 |
| ENSG00000005194 | CIAPIN1 | CIAPIN1 | cytokine induced apoptosis inhibitor 1 | 16q21 |
| ENSG00000165055.16 | METTL2B | METTL2B | methyltransferase 2B, methylcytidine | 7q32.1 |
| ENSG00000164169.13 | PRMT10 | PRMT9 | protein arginine methyltransferase 9 | 4q31.23 |
| ENSG00000150756.14 | FAM173B | ATPSCKMT | ATP synthase c subunit lysine N-methyltransferase | 5p15.2 |
| ENSG00000170439.8 | METTL7B | TMT1B | thiol methyltransferase 1B | 12q13.2 |
| ENSG00000146834.15 | MEPCE | MEPCE | methylphosphate capping enzyme | 7q22.1 |
| ENSG00000003509.16 | C2orf56 | NDUFAF7 | NADH:ubiquinone oxidoreductase complex assembly factor 7 | 2p22.2 |
| ENSG00000121486.12 | TRM1L | TRMT1L | tRNA methyltransferase 1 like | 1q25.3 |
| ENSG00000066651.20 | TRMT11 | TRMT11 | tRNA methyltransferase 11 homolog | 6q22.32 |
| ENSG00000186666.6 | BCDIN3D | BCDIN3D | BCDIN3 domain containing RNA methyltransferase | 12q13.12 |
| ENSG00000143919.15 | C2orf34 | CAMKMT | calmodulin-lysine N-methyltransferase | 2p21 |
| ENSG00000165644.11 | COMTD1 | COMTD1 | catechol-O-methyltransferase domain containing 1 | 10q22.2 |
| ENSG00000127804.13 | METT10D | METTL16 | methyltransferase 16, N6-methyladenosine | 17p13.3 |

| | | | | |
|---|---|---|---|---|
| ENSG00000142453.12 | CARM1 | CARM1 | coactivator associated arginine methyltransferase 1 | 19p13.2 |
| ENSG00000181038.14 | C17orf95 | METTL23 | methyltransferase like 23 | 17q25.2 |
| ENSG00000139160.13 | C12orf72 | ETFBKMT | electron transfer flavoprotein subunit beta lysine methyltransferase | 12p11.21 |
| ENSG00000108592.17 | FTSJ3 | FTSJ3 | FtsJ RNA 2'-O-methyltransferase 3 | 17q23.3 |
| ENSG00000155275.19 | C4orf23 | TRMT44 | tRNA methyltransferase 44 homolog | 4p16.1 |
| ENSG00000180917.18 | FTSJD1 | CMTR2 | cap methyltransferase 2 | 16q22.2 |
| ENSG00000099899.15 | TRMT2A | TRMT2A | tRNA methyltransferase 2 homolog A | 22q11.21 |
| ENSG00000137200.13 | FTSJD2 | CMTR1 | cap methyltransferase 1 | 6p21.2 |
| ENSG00000156017.13 | C9orf41 | CARNMT1 | carnosine N-methyltransferase 1 | 9q21.13 |
| ENSG00000165171.11 | WBSCR27 | METTL27 | methyltransferase like 27 | 7q11.23 |
| ENSG00000127720.8 | C12orf26 | METTL25 | methyltransferase like 25 | 12q21.31 |
| ENSG00000010165.20 | METTL13 | METTL13 | methyltransferase 13, eEF1A lysine and N-terminal methyltransferase | 1q24.3 |
| ENSG00000186523.15 | FAM86B1 | FAM86B1 | family with sequence similarity 86 member B1 | 8p23.1 |
| ENSG00000179299.17 | NSUN7 | NSUN7 | NOP2/Sun RNA methyltransferase family member 7 | 4p14 |
| ENSG00000138780.15 | GSTCD | GSTCD | glutathione S-transferase C-terminal domain containing | 4q24 |
| ENSG00000206562.12 | METTL6 | METTL6 | methyltransferase 6, methylcytidine | 3p25.1 |
| ENSG00000241058.4 | NSUN6 | NSUN6 | NOP2/Sun RNA methyltransferase 6 | 10p12.31 |
| ENSG00000104885.19 | DOT1L | DOT1L | DOT1 like histone lysine methyltransferase | 19p13.3 |
| ENSG00000029639.11 | TFB1M | TFB1M | transcription factor B1, mitochondrial | 6q25.3 |
| ENSG00000144401.14 | FAM119A | METTL21A | methyltransferase 21A, HSPA lysine | 2q33.3 |
| ENSG00000123427.17 | FAM119B | EEF1AKMT3 | EEF1A lysine methyltransferase 3 | 12q14.1 |
| ENSG00000137760.15 | ALKBH8 | ALKBH8 | alkB homolog 8, tRNA methyltransferase | 11q22.3 |
| ENSG00000117481.11 | NSUN4 | NSUN4 | NOP2/Sun RNA methyltransferase 4 | 1p33 |
| ENSG00000143303.12 | C1orf66 | METTL25B | methyltransferase like 25B | 1q23.1 |
| ENSG00000166166.13 | TRMT61A | TRMT61A | tRNA methyltransferase 61A | 14q32.33 |
| ENSG00000118894.15 | FAM86A | EEF2KMT | eukaryotic elongation factor 2 lysine methyltransferase | 16p13.3 |
| ENSG00000188917.15 | TRMT2B | TRMT2B | tRNA methyltransferase 2 homolog B | Xq22.1 |
| ENSG00000087995.16 | METTL2A | METTL2A | methyltransferase 2A, methylcytidine | 17q23.2 |

| | | | | |
|---|---|---|---|---|
| ENSG00000198890.9 | PRMT6 | PRMT6 | protein arginine methyltransferase 6 | 1p13.3 |
| ENSG00000168300.14 | PCMTD1 | PCMTD1 | protein-L-isoaspartate (D-aspartate) O-methyltransferase domain containing 1 | 8q11.23 |
| ENSG00000130305.17 | NSUN5 | NSUN5 | NOP2/Sun RNA methyltransferase 5 | 7q11.23 |
| ENSG00000137574.11 | TGS1 | TGS1 | trimethylguanosine synthase 1 | 8q12.1 |
| ENSG00000130731.16 | C16orf13 | METTL26 | methyltransferase like 26 | 16p13.3 |
| ENSG00000126457.22 | PRMT1 | PRMT1 | protein arginine methyltransferase 1 | 19q13.33 |
| ENSG00000103254.10 | FAM173A | ANTKMT | adenine nucleotide translocase lysine methyltransferase | 16p13.3 |
| ENSG00000137364 | TPMT | TPMT | thiopurine S-methyltransferase | 6p22.3 |
| ENSG00000138050.15 | THUMPD2 | THUMPD2 | THUMP domain containing 2 | 2p22.1 |
| ENSG00000067365.15 | C16orf68 | METTL22 | methyltransferase 22, Kin17 lysine | 16p13.2 |
| ENSG00000134077.16 | THUMPD3 | THUMPD3 | THUMP domain containing 3 | 3p25.3 |
| ENSG00000148335.15 | METTL11A | NTMT1 | N-terminal Xaa-Pro-Lys N-methyltransferase 1 | 9q34.11 |
| ENSG00000171103.11 | TRMT61B | TRMT61B | tRNA methyltransferase 61B | 2p23.2 |
| ENSG00000197006.15 | METTL9 | METTL9 | methyltransferase like 9 | 16p12.2 |
| ENSG00000162851.8 | TFB2M | TFB2M | transcription factor B2, mitochondrial | 1q44 |
| ENSG00000178694.10 | NSUN3 | NSUN3 | NOP2/Sun RNA methyltransferase 3 | 3q11.2 |
| ENSG00000165792.18 | METT11D1 | METTL17 | methyltransferase like 17 | 14q11.2 |
| ENSG00000123600.21 | METTL8 | METTL8 | methyltransferase 8, methylcytidine | 2q31.1 |
| ENSG00000100483.14 | C14orf138 | VCPKMT | valosin containing protein lysine methyltransferase | 14q21.3 |
| ENSG00000185432.12 | METTL7A | TMT1A | thiol methyltransferase 1A | 12q13.12 |
| ENSG00000214435.9 | AS3MT | AS3MT | arsenite methyltransferase | 10q24.32 |
| ENSG00000111218.12 | PRMT8 | PRMT8 | protein arginine methyltransferase 8 | 12p13.32 |
| ENSG00000138382.15 | METTL5 | METTL5 | methyltransferase 5, N6-adenosine | 2q31.1 |
| ENSG00000122435.11 | CCDC76 | TRMT13 | tRNA methyltransferase 13 homolog | 1p21.2 |
| ENSG00000203880.12 | PCMTD2 | PCMTD2 | protein-L-isoaspartate (D-aspartate) O-methyltransferase domain containing 2 | 20q13.33 |
| ENSG00000132600.18 | PRMT7 | PRMT7 | protein arginine methyltransferase 7 | 16q22.1 |
| ENSG00000104907.13 | TRMT1 | TRMT1 | tRNA methyltransferase 1 | 19p13.13 |

| | | | | |
|---|---|---|---|---|
| ENSG00000132423.12 | COQ3 | COQ3 | coenzyme Q3, methyltransferase | 6q16.2 |
| ENSG00000250305.9 | C8orf79 | TRMT9B | tRNA methyltransferase 9B (putative) | 8p22 |
| ENSG00000037897.17 | METTL1 | METTL1 | methyltransferase 1, tRNA methylguanosine | 12q14.1 |
| ENSG00000068438.15 | FTSJ1 | FTSJ1 | FtsJ RNA 2'-O-methyltransferase 1 | Xp11.23 |
| ENSG00000122687.19 | FTSJ2 | MRM2 | mitochondrial rRNA methyltransferase 2 | 7p22.3 |
| ENSG00000205629.12 | LCMT1 | LCMT1 | leucine carboxyl methyltransferase 1 | 16p12.1 |
| ENSG00000086189.11 | DIMT1L | DIMT1 | DIM1 rRNA methyltransferase and ribosome maturation factor | 5q12.1 |
| ENSG00000156239.12 | N6AMT1 | N6AMT1 | N-6 adenine-specific DNA methyltransferase 1 | 21q21.3 |
| ENSG00000114735.10 | HEMK1 | HEMK1 | HemK methyltransferase family member 1 | 3p21.31 |
| ENSG00000150456.11 | N6AMT2 | EEF1AKMT1 | EEF1A lysine methyltransferase 1 | 13q12.11 |
| ENSG00000168228.16 | ZCCHC4 | ZCCHC4 | zinc finger CCHC-type containing 4 | 4p15.2 |
| ENSG00000165819.12 | METTL3 | METTL3 | methyltransferase 3, N6-adenosine-methyltransferase complex catalytic subunit | 14q11.2 |
| ENSG00000101574.15 | METTL4 | METTL4 | methyltransferase 4, N6-adenosine | 18p11.32 |
| ENSG00000145388.15 | METTL14 | METTL14 | methyltransferase 14, N6-adenosine-methyltransferase subunit | 4q26 |
| ENSG00000085276.19 | MECOM | MECOM | MDS1 and EVI1 complex locus | 3q26.2 |
| ENSG00000167548.18 | MLL2 | KMT2D | lysine methyltransferase 2D | 12q13.12 |
| ENSG00000099381.19 | SETD1A | SETD1A | SET domain containing 1A, histone lysine methyltransferase | 16p11.2 |
| ENSG00000101945.17 | SUV39H1 | SUV39H1 | SUV39H1 histone lysine methyltransferase | Xp11.23 |
| ENSG00000057657.17 | PRDM1 | PRDM1 | PR/SET domain 1 | 6q21 |
| ENSG00000109685.19 | WHSC1 | NSD2 | nuclear receptor binding SET domain protein 2 | 4p16.3 |
| ENSG00000141956.14 | PRDM15 | PRDM15 | PR/SET domain 15 | 21q22.3 |
| ENSG00000118058.24 | MLL | KMT2A | lysine methyltransferase 2A | 11q23.3 |
| ENSG00000116731.23 | PRDM2 | PRDM2 | PR/SET domain 2 | 1p36.21 |
| ENSG00000085276.19 | MECOM | TTLL12 | tubulin tyrosine ligase like 12 | 22q13.2 |
| ENSG00000100304.13 | TTLL12 | SETDB1 | SET domain bifurcated histone lysine methyltransferase 1 | 1q21.3 |
| ENSG00000143379.13 | SETDB1 | EZH2 | enhancer of zeste 2 polycomb repressive complex 2 subunit | 7q36.1 |
| ENSG00000106462.12 | EZH2 | KMT5B | lysine methyltransferase 5B | 11q13.2 |
| ENSG00000110066.15 | SUV420H1 | SETMAR | SET domain and mariner transposase fusion gene | 3p26.1 |

| | | | | |
|---|---|---|---|---|
| ENSG00000170364.13 | SETMAR | SMYD5 | SMYD family member 5 | 2p13.2 |
| ENSG00000135632.12 | SMYD5 | SETD3 | SET domain containing 3, actin histidine methyltransferase | 14q32.2 |
| ENSG00000183576.13 | SETD3 | KMT5C | lysine methyltransferase 5C | 19q13.42 |
| ENSG00000133247.14 | SUV420H2 | ZFPM1 | zinc finger protein, FOG family member 1 | 16q24.2 |
| ENSG00000179588.9 | ZFPM1 | SMYD4 | SET and MYND domain containing 4 | 17p13.3 |
| ENSG00000186532.12 | SMYD4 | KMT2E | lysine methyltransferase 2E (inactive) | 7q22.3 |
| ENSG00000005483.22 | MLL5 | SMYD1 | SET and MYND domain containing 1 | 2p11.2 |
| ENSG00000115593.15 | SMYD1 | SETD9 | SET domain containing 9 | 5q11.2 |
| ENSG00000155542.12 | C5orf35 | KMT2C | lysine methyltransferase 2C | 7q36.1 |
| ENSG00000055609.21 | MLL3 | SETD6 | SET domain containing 6, protein lysine methyltransferase | 16q21 |
| ENSG00000103037.12 | SETD6 | SETD7 | SET domain containing 7, histone lysine methyltransferase | 4q31.1 |
| ENSG00000145391.14 | SETD7 | ZFPM2 | zinc finger protein, FOG family member 2 | 8q23 |
| ENSG00000169946.14 | ZFPM2 | EZH1 | enhancer of zeste 1 polycomb repressive complex 2 subunit | 17q21.2 |
| ENSG00000108799.13 | EZH1 | EZH2 | enhancer of zeste 2 polycomb repressive complex 2 subunit | 7q36.1 |
| ENSG00000204371 | EHMT2 | EHMT2 | euchromatic histone lysine methyltransferase 2 | 6p21.33 |
| ENSG00000165671.22 | NSD1 | NSD1 | nuclear receptor binding SET domain protein 1 | 5q35.3 |
| ENSG00000136169.17 | SETDB2 | SETDB2 | SET domain bifurcated histone lysine methyltransferase 2 | 13q14.2 |
| ENSG00000181555.22 | SETD2 | SETD2 | SET domain containing 2, histone lysine methyltransferase | 3p21.31 |
| ENSG00000147548.17 | WHSC1L1 | NSD3 | nuclear receptor binding SET domain protein 3 | 8p11.23 |
| ENSG00000168137.20 | SETD5 | SETD5 | SET domain containing 5 | 3p25.3 |
| ENSG00000147596.4 | PRDM14 | PRDM14 | PR/SET domain 14 | 8q13.3 |
| ENSG00000112238.12 | PRDM13 | PRDM13 | PR/SET domain 13 | 6q16.2 |
| ENSG00000130711.5 | PRDM12 | PRDM12 | PR/SET domain 12 | 9q34.12 |
| ENSG00000152455.16 | SUV39H2 | SUV39H2 | SUV39H2 histone lysine methyltransferase | 10p13 |
| ENSG00000185420.19 | SMYD3 | SMYD3 | SET and MYND domain containing 3 | 1q44 |
| ENSG00000181090.21 | EHMT1 | EHMT1 | euchromatic histone lysine methyltransferase 1 | 9q34.3 |
| ENSG00000175213.3 | ZNF408 | ZNF408 | zinc finger protein 408 | 11p11.2 |
| ENSG00000142611.17 | PRDM16 | PRDM16 | PR/SET domain 16 | 1p36.32 |

| | | | | |
|---|---|---|---|---|
| ENSG00000183955.14 | SETD8 | KMT5A | lysine methyltransferase 5A | 12q24.31 |
| ENSG00000019485.14 | PRDM11 | PRDM11 | PR/SET domain 11 | 11p11.2 |
| ENSG00000170325.16 | PRDM10 | PRDM10 | PR/SET domain 10 | 11q24.3 |
| ENSG00000164256.11 | PRDM9 | PRDM9 | PR/SET domain 9 | 5p14.2 |
| ENSG00000152784.16 | PRDM8 | PRDM8 | PR/SET domain 8 | 4q21.21 |
| ENSG00000126856.16 | PRDM7 | PRDM7 | PR/SET domain 7 | 16q24.3 |
| ENSG00000061455.11 | PRDM6 | PRDM6 | PR/SET domain 6 | 5q23.2 |
| ENSG00000138738.11 | PRDM5 | PRDM5 | PR/SET domain 5 | 4q27 |
| ENSG00000116539.14 | ASH1L | ASH1L | ASH1 like histone lysine methyltransferase | 1q22 |
| ENSG00000143499.14 | SMYD2 | SMYD2 | SET and MYND domain containing 2 | 1q32.3 |
| ENSG00000185917.14 | SETD4 | SETD4 | SET domain containing 4 | 21q22.12 |
| ENSG00000110851.12 | PRDM4 | PRDM4 | PR/SET domain 4 | 12q23.3 |
| ENSG00000272333.8 | WBP7 | KMT2B | lysine methyltransferase 2B | 19q13.12 |
| ENSG00000139718.12 | SETD1B | SETD1B | SET domain containing 1B, histone lysine methyltransferase | 12q24.31 |
| ENSG00000059588.10 | TARBP1 | TARBP1 | TAR (HIV-1) RNA binding protein 1 | 1q42.2 |
| ENSG00000198917.13 | C9orf114 | SPOUT1 | SPOUT domain containing methyltransferase 1 | 9q34.11 |
| ENSG00000278619 | MRM1 | MRM1 | mitochondrial rRNA methyltransferase 1 | 17q12 |
| ENSG00000165275.10 | RG9MTD3 | TRMT10B | tRNA methyltransferase 10B | 9p13.2 |
| ENSG00000174173.7 | RG9MTD1 | TRMT10C | tRNA methyltransferase 10C, mitochondrial RNase P subunit | 3q12.3 |
| ENSG00000145331.14 | RG9MTD2 | TRMT10A | tRNA methyltransferase 10A | 4q23 |
| ENSG00000126749.16 | EMG1 | EMG1 | EMG1 N1-specific pseudouridine methyltransferase | 12p13.31 |
| ENSG00000171861.11 | RNMTL1 | MRM3 | mitochondrial rRNA methyltransferase 3 | 17p13.3 |
| ENSG00000162623.16 | TYW3 | TYW3 | tRNA-yW synthesizing protein 3 homolog | 1p31.1 |
| ENSG00000145692.15 | BHMT | BHMT | betaine--homocysteine S-methyltransferase | 5q14.1 |
| ENSG00000132840.10 | BHMT2 | BHMT2 | betaine--homocysteine S-methyltransferase 2 | 5q14.1 |
| ENSG00000116237.16 | ICMT | ICMT | isoprenylcysteine carboxyl methyltransferase | 1p36.31 |
| ENSG00000137404 | NRM | NRM | nurim | 6p21.33 |
| ENSG00000133027.18 | PEMT | PEMT | phosphatidylethanolamine N-methyltransferase | 17p11.2 |

| | | | | |
|---|---|---|---|---|
| ENSG00000116984.15 | MTR | MTR | 5-methyltetrahydrofolate-homocysteine methyltransferase | 1q43 |
| ENSG00000117543 | DPH5 | DPH5 | diphthamide biosynthesis 5 | 1p21.2 |
| ENSG00000145996.11 | CDKAL1 | CDKAL1 | CDK5 regulatory subunit associated protein 1 like 1 | 6p22.3 |
| ENSG00000101391.21 | CDK5RAP1 | CDK5RAP1 | CDK5 regulatory subunit associated protein 1 | 20q11.21 |
| ENSG00000134014.17 | ELP3 | ELP3 | elongator acetyltransferase complex subunit 3 | 8p21.1 |
| ENSG00000136444.10 | RSAD1 | RSAD1 | radical S-adenosyl methionine domain containing 1 | 17q21.33 |

Supplementary Table 1. Identified methyl transferase genes used for analysis of RNA sequencing data.
